# Emergence of single cell mechanical behavior and polarity within epithelial monolayers drives collective cell migration

**DOI:** 10.1101/2019.12.16.875567

**Authors:** Shreyansh Jain, Victoire M.L. Cachoux, Gautham H.N.S. Narayana, Simon de Beco, Joseph D’Alessandro, Victor Cellerin, Tianchi Chen, Mélina L. Heuzé, Philippe Marcq, René-Marc Mège, Alexandre J. Kabla, Chwee Teck Lim, Benoit Ladoux

## Abstract

The directed migration of cell collectives is essential in various physiological processes, such as epiboly, intestinal epithelial turnover, and convergent extension during morphogenesis as well as during pathological events like wound healing and cancer metastasis^1,2^. Collective cell migration leads to the emergence of coordinated movements over multiple cells. Our current understanding emphasizes that these movements are mainly driven by large-scale transmission of signals through adherens junctions^3,4^. In this study, we show that collective movements of epithelial cells can be triggered by polarity signals at the single cell level through the establishment of coordinated lamellipodial protrusions. We designed a minimalistic model system to generate one-dimensional epithelial trains confined in ring shaped patterns that recapitulate rotational movements observed *in vitro* in cellular monolayers and *in vivo* in genitalia or follicular cell rotation^5–7^. Using our system, we demonstrated that cells follow coordinated rotational movements after the establishment of directed Rac1-dependent polarity over the entire monolayer. Our experimental and numerical approaches show that the maintenance of coordinated migration requires the acquisition of a front-back polarity within each single cell but does not require the maintenance of cell-cell junctions. Taken together, these unexpected findings demonstrate that collective cell dynamics in closed environments as observed in multiple *in vitro* and *in vivo* situations^5,6,8,9^ can arise from single cell behavior through a sustained memory of cell polarity.

## Main Text

The ability of cells to migrate collectively is crucial in shaping organisms during the complex morphogenetic events of development, and for several physiological and pathological events like wound healing and cancer metastasis^1,2^. Single cell migration is associated with a front-back polarity that includes the formation of a lamellipodial structure at the leading edge^10,11^. Even though this mode of migration is still under intense research^12^, it is now clearly established that the protrusive activity driven by actin polymerization at the cell front leads to forward movement in a directional and persistent fashion^13,14^. Collective movements require a higher degree of complexity and are thus less well understood. Collectively migrating cells display a complex range of front-rear polarization and mechanical coupling behaviors that depend on their position within the migrating monolayer^15,16^. Collective migration behaviors occur under a broad range of external constraints that induce the appearance of highly motile mesenchymal-like ‘leader’ cells^17^, the local guidance of small cohorts of ‘follower’ cells^18^, and large-scale movements within the bulk of cell monolayers^19^. The emergence of these polarized cellular assemblies thus results from the integration of intra- and extra-cellular biomechanical cues that cooperate to steer and maintain the migration of cohesive groups^20^.

Both *in vivo* and *in vitro* studies on collective cell movements have uncovered the importance of front-rear polarization at the single cell level, for instance through the formation of “cryptic” lamellipodia^21,22^, but also that of a large-scale coupling between cells through intercellular adhesions^3,4,23,24^. Moreover, studies *in vivo* on Drosophila follicular cells and genitalia cells, as well as *in vitro* cell migration systems show that cell monolayers can coordinate and rotate persistently even without free boundaries^5,7,8^. Interestingly, the emergence of collective cell migration in *Drosophila* follicular cells or more recently in mammalian intestinal turnover shows front-back polarity patterns at the single cell level^9^. Overall, the mechanisms behind epithelial cell migration remain largely unknown and controversial.

Using a micro-contact printing method on polydimethyl-siloxane (PDMS), we confined trains of Madin-Darby canine kidney epithelial cells (MDCK) on fibronectin-coated annular rings of controlled geometrical dimensions^25^. We mainly performed experiments using rings of 200 µm in outer diameter and 20 µm in width to limit cell migration along the ring to a one-dimensional track (Fig. 1a). This configuration minimizes lateral intercellular interactions and thus simplifies the analysis of the experimental system. After cell seeding, cells coalesced and spread to form random-sized short trains (Fig. 1b). We tracked the coordination of these trains by quantifying the coordination parameter *D*, defined as the average of the sum of cross products between cell velocity unit vector (v) measured from particle image velocimetry (PIV) and its respective position vector from the center of the ring 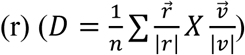(n, total number of points in a given space) such that the normalized values of +1, −1, or 0 indicated clockwise, anticlockwise, or no preferential direction, respectively. At sub-confluency, cell trains displayed oscillatory migratory behaviors with no preferential direction (**Movie 1**) (Fig. 1c, d **before 12h**). However, we observed a surge in coordination parameter and average velocity as cell trains gradually met and fused into larger trains (Fig. 1b, c, **d after 12 h**). Eventually the coordination parameter came to a final, lasting plateau at ±1 when; cells reached confluency and began to rotate persistently in either a clockwise or anticlockwise direction with equal probability (Fig. 1d, **at 20 h**) (**Extended Data Fig. 1a**) (**Movie 1**). The coordination parameter gradually decreased in the end due to continued cell proliferation, a property reminiscent of cell jamming^7,8,26,27^(**Extended Data Fig. 1b**). When cells were treated with mitomycin to maintain a constant cell number, collective cell behaviors were unaffected over the first 40 hours, which was enough to reach confluency (**Extended Data Fig. 1b**). The average speed of rotation was higher for cells under mitomycin treatment (**Extended Data Fig. 1c,d**), probably due to the flow disturbance caused by continuing cell division under normal conditions^28^. To generalize our findings, we first varied the geometrical parameters characterizing cell confinement, including widths, perimeters and curvatures of closed patterns. Under these conditions, we did not affect the global rotational movements, demonstrating that this collective cell behavior was qualitatively independent of geometrical constraints (Fig. 1e,f) (**Movie 2**). In addition, we showed that our findings could be extended to other epithelial cell lines including Caco2 (Human epithelial colorectal adenocarcinoma cells) and Eph4 (Mouse mammary gland epithelial cells) that displayed similar behaviors (**Extended Data Fig. 1e**) (**Movie 3**).

**Figure. 1.**
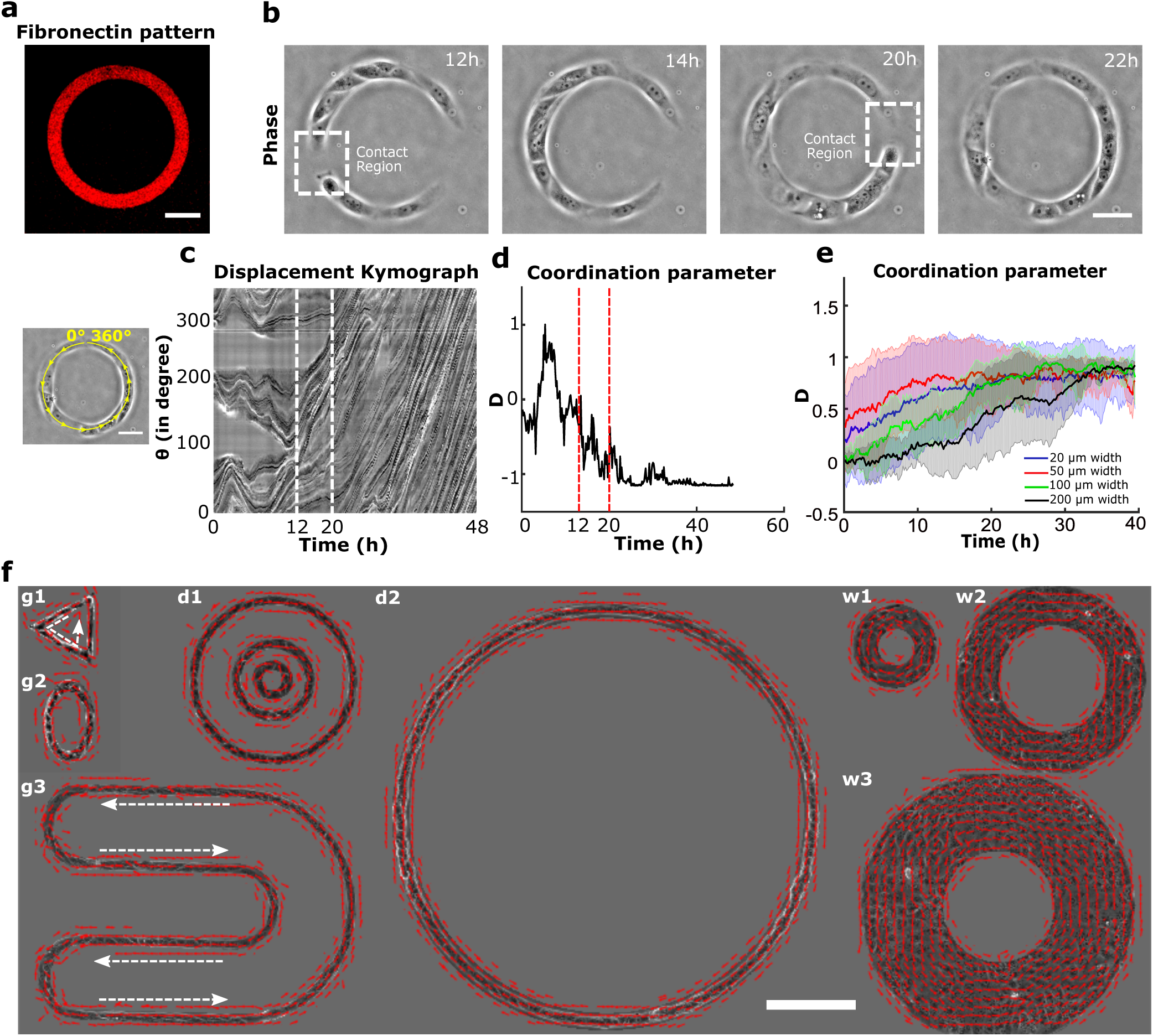
Experimental setup and directed collective migration of MDCK trains. **a**, Micro-contact printed pattern of Cy3 conjugated fibronectin in the shape of annular ring (ring outer diameter = 200 µm, track width = 20 µm). **b**, Phase contrast images of train of MDCK cells after initial seeding and spreading. Dashed white box indicates the contact regions where small cell trains merged into larger trains at12h and 20h. **c**, Spatio-temporal displacement of cells traced along the circumferential midline of the ring track. Small train merging event is indicated by red dashed line. **d**, Evolution of coordination parameter, *D*, of cell trains in the ring with time (at t=20h cell train in ring begins to rotate in anti-clockwise direction). **e**, Coordination parameter, *D*, of cells in ring with different width: 20 µm (n=62, m=3),50 µm (n=27, m=2),100 µm (n=8), 200 µm (n=8) respectively, show a collective migration of cell trains independent of track width. **f**, Velocity vectors representing directed collective cell migrations for a variety of confined geometries (g1-triangle, g2-ellipse, g3-U shape), diameter (d1:outer-400 µm, middle-200 µm, inner-100 µm, d2-1mm), and width (w1-50 µm, w2-100 µm, w3-200 µm). Scale bars a, b, c: 50 µm and f: 200 µm.

We first observed a spontaneous process of symmetry breaking that initiated global migration into one preferential direction (Fig. 2a, **20 min**). The last contact between two leading cells that extend lamellipodial protrusions led to the change in direction of only one of the cells, a mechanism that may be partly reminiscent of contact inhibition of locomotion (CIL)^29^. Since CIL is based on opposite repolarization of both cells away from their contact site, we sought to study the dynamics of front-rear cell polarization in the ring using a fluorescent biosensor (P21 activated kinase Binding Protein, PBD) of active Rac1 and Cdc42^30^. We observed a reorientation of the PBD biosensor gradient in one of the two leading cells upon contact (Fig. 2a, **40 min**) (**Movie 4**) with time scales of cell repolarization as fast as 3~4 min (**Extended Data Fig. 2a,b**).

**Figure 2.**
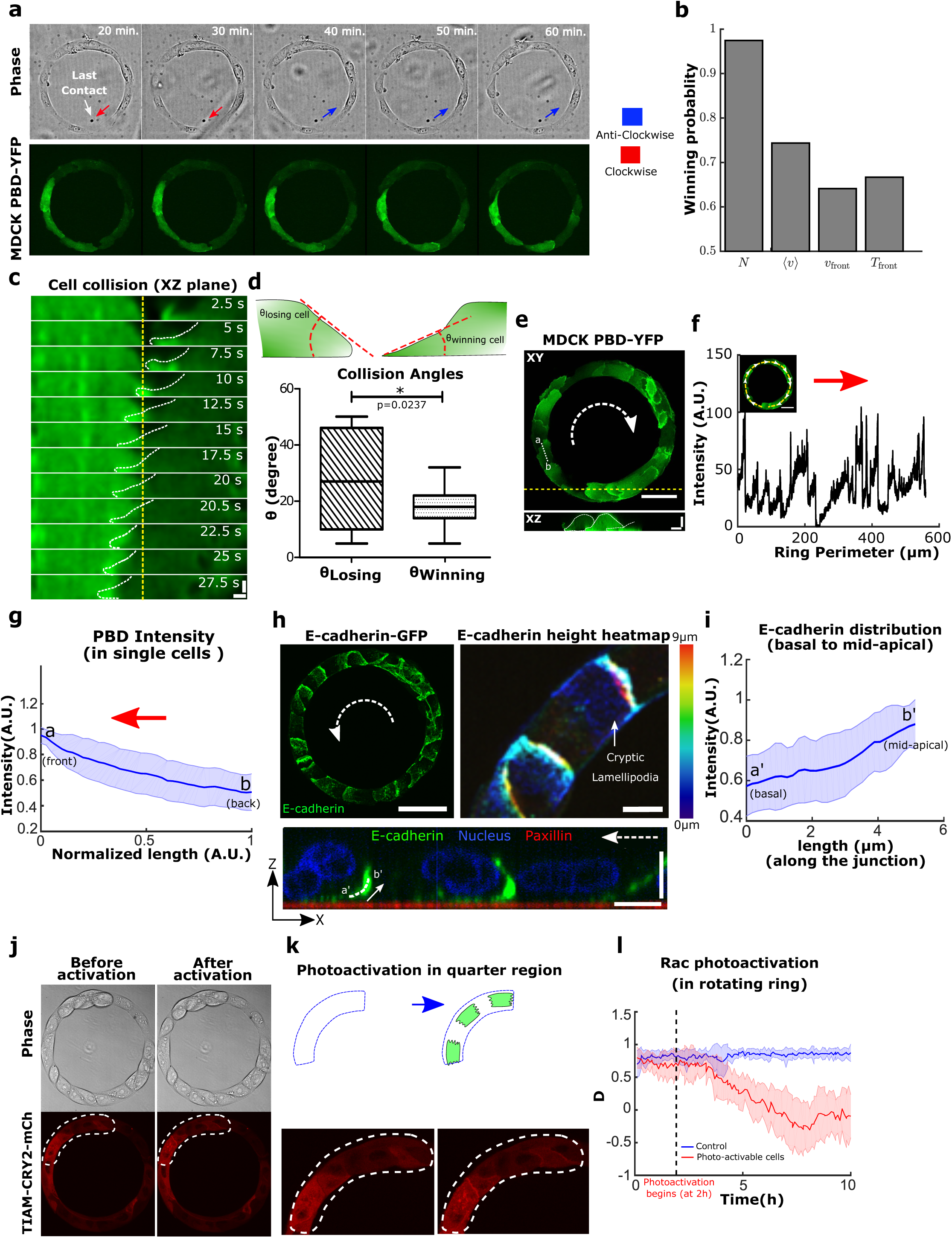
Symmetry breaking in the epithelial train leads to the emergence of single cell polarity in migrating epithelia. **a**, Confocal images (basal plane) of MDCK-PBD-YFP cells just before and after the establishment of last contact in the cell train. Phase contrast images show two oppositely migrating cell trains (red and blue arrows indicate the clockwise and anticlockwise direction respectively). Region indicating the establishment of last contact before confluency marked by solid circle (at 30 min.); Confocal sequential images showing the intensity of PBD-YFP biosensor signal during the last colliding event (dashed circle, at 40 min). **b**, Winning probability plot of cell trains upon collision, based on different parameters *N* (the train with large number of cells)*, <v>* (average speeds), *V_front_* (front velocities), *T_front_* (traction forces at the colliding front) (n=39, m=3). These parameters of colliding trains were measured 3 timepoints (i.e. 30 min.) before collision. **c**, Lamellipodial contact dynamics of the colliding cells as observed in XZ section of MDCK PBD-YFP cells (XZ Scale bar: 2 µm). **d**, Winning and losing cell angles (schematic) as measured by the XZ section of the colliding cells (n=11, m=4). **e**, Confocal image of basal plane in MDCK-PBD cells showing the PBD biosensor distribution in cells. Dashed arrow indicates the rotation direction of the ring. XZ plane shows the establishment of Rac1 rich cryptic lamellipodia’s in cells (Scale bar: 4 µm). **f**, PBD intensity profile traced along the circumferential midline of ring track shown in **e**. **g**, Normalized PBD intensity profile in single cells (along line -ab in panel **e**, n=102 from 10 rings, m=3). The red arrow indicates the direction of cell migration. **h**, (Left) Confocal immunofluorescence image of E-cadherin (maximum intensity projection). Dashed white arrow indicates the direction of rotation in the ring. (Right) Expanded view (dashed yellow box in **h**, Scale bar: 10 µm) shows E-cadherin distribution height heatmap. ZX plane (bottom) corresponding to the dashed yellow region shows the formation of cryptic lamellipodia underneath the neighboring cells (Scale bar: 10 µm). Dashed white arrow indicates the direction of migration. **i**, Intensity profile of E-cadherin along the basal to mid-apical region (n=40 from 7 rings, m=1) (a’-b’, marked in ZX plane of **h**, scale bar: 10 µm). **j-k**, Optogenetic-based homogeneous relocalization of TIAM (see methods) at cell membranes in a quarter of the ring (marked by dashed line). The schematic indicates the polarity perturbation post photoactivation. **l**, Evolution of the coordination parameter *D* as a function of time after photoactivation. Photo-activable cells in red (n=11, m=2) and WT control cells after similar photoactivation in blue (n=9, m=1). All scale bars unless mentioned specifically: 50 µm.

To further investigate the principles leading to this repolarization, we tested different parameters of both trains just before the last collision (3 time points before, i.e. 30 minutes). Our results indicated that the final direction of rotation at confluency was best predicted by the number of cells constituting the last colliding trains, then by their front speeds, with the winning train being the longest and the fastest (Fig.2b). Since prominent leader cells lead to high levels of orientation and polarization of followers^31^, we reasoned that winning cells should be then evidenced by larger lamellipodial protrusions. We thus analyzed the local protrusive activity of colliding cells before confluency. By measuring the contact angles of these two cells, we indeed found that the most spread cell with smaller angle of contact (i.e. large spread of lamellipodia) won over the other which repolarized in the opposite direction. Even more, the winning cell extended its lamellipodia beneath the other one eventually changing its polarity (Fig. 2c,d,e). Altogether, our findings demonstrated that coordinated rotational motion was driven by the repolarization of the smaller moving cell groups upon frontal collision due to local biased protrusive activity of leading cells.

Shortly after reaching confluency, cells exhibited a persistent collective rotational movement. We then studied the maintenance of this persistent coordination over long-time scales. We first followed the distribution of the PBD sensor as an indicator of polarity gradient of Rac1 (**Movie 5**) (Fig. 2e,f,g). Surprisingly, we found that cells established a unidirectional polarity gradient at single cell level within collectively migrating monolayers. This observation was not restricted to 1-D confined monolayers but was also present in epithelial cells presenting lateral contacts as observed in wider rotating rings (Extended Data Fig. 2c,d). For the sake of simplicity, we thus focused our analysis on 1-D configurations. We then reasoned that the emergence of single cell polarization should display antagonistic activation of RhoA and Rac1 signals. Analyzing the active RhoA localization in cells using AHPH domain sensor^32^ further demonstrated the presence of an active contractility gradient at a single cell level opposite to the PBD gradient (**Extended Data Fig. 2e,f,g,h,i**). Such patterns of Rho GTPases activity were consistent with a picture of individual cell behavior controlling the movement of cell groups, rather than supracellular contractility as observed upon migration with free boundaries^33^. To further investigate the organization of these clusters, we performed immunostaining of cell-substrate and cell-cell adhesions based on paxillin and E-cadherin staining. It confirmed the presence of cryptic lamellipodia underneath the adjacent cell (Fig. 2h, **Z-X plane**; **Extended Data Fig. 3a, b, c, d, e, f**). Cadherin distribution also revealed a dissymmetrical organization with a gradient from basal to mid-apical level of the cells, including diffused clusters on the dorsal surface of the cryptic lamellipodium and a more concentrated E-cadherin region at the mid-apical region of the cells (Fig. 2i). We next sought to further understand the long-term persistency of these collective movements. The assembly of cryptic lamellipodia underneath the adjacent cell body led to the confinement of the protrusive structures. Cellular confinement by itself can promote actin retrograde flows and cell polarity^34^ that in turn reinforce cell persistency^14^. By introducing artificial confinement, we could demonstrate that confined lamellipodia promoted the assembly of large protrusive structures with higher stability and extension speeds, which could favor increased persistent cell migration as a collective (**Extended Data Fig. 4a,b,c,d**). This finding was even strengthened by the observation of a preferential alignment of the centrosome in front of the nucleus towards the direction of collective motion (around 60~70% of cells), which was not observed before rotation (**Extended Data Fig. 5a,b**).

To test the principles behind the maintenance of persistent coordinated migration, we perturbed the system with the Rac-1 inhibitor Z62954982 (100 µM). It induced a rapid disappearance of PBD sensor distribution gradients in cells, and blocked rotation (**Extended Data Fig. 6a,b,c,d,e**). We then perturbed actin polymerization by preventing the branching of filaments through Arp2/3 inhibition by CK-666 (100 µM) (**Extended Data Fig. 6f**). Similarly, we observed a gradual decrease of cell velocities followed by a complete arrest of rotational motion. It suggests that both actin polymerization and lamellipodium formation are crucial to maintain such collective movements. We then examined the effect of local perturbations in front-back polarity of a few cells on the collective coordination of the entire system. To do so, we used optogenetics to flatten-out the Rac1 gradients in targeted groups of a few cells in each ring and thus perturb their polarity (**Movie 6**). This homogeneous whole cell Rac1 activation led to random extensions of lamellipodia among the targeted subgroups of cells, thereby stopping their directed migration. This progressively led to a total inhibition of the rotational migration over the entire ring (Fig. 2j,k,l and **Extended Data Fig. 7a,b**). These results pointed towards a process of cellular self-organization through polar extension of lamellipodia at the single cell level. However, other mechanisms could have contributed to such collective movements including propagating waves through acto-myosin contractility^24,35^. We thus analyzed the impact of Myosin II inhibition upon a high Blebbistatin concentration (80 µM). To our surprise, the coordination parameter dropped down gradually to reach half of its maximal value over several hours, showing myosin II contractility had less severe impact on collective behavior than Rac1 and Arp2/3 inhibitions (**Extended Data 6f**).

Our findings revealed that the initiation of coordinated movements occurred when cells reached confluency after the last collision of two migrating trains and unveiled the crucial role of lamellipodial protrusions and single cell polarity. We thus further investigated whether purely physical interactions by volume exclusion in a packed system could initiate coordinated rotational movement at confluent density or whether cell-cell adhesion was involved for initiating and maintaining a persistent collective rotation in the absence of a front edge. To do so, we perturbed cadherin-based adhesions by knocking down α-catenin^36^. Unlike control cells, α-catenin deficient cells did not initiate large-scale coordinated movements. Similar results were obtained with cells expressing the transcription factor Snail-1 in which E-cadherin mediated adhesion was also impaired^7^, demonstrating that cell-cell junctions were required to initiate collective movements (Extended Data Fig. 1f) (**Movie 7**).

We then analyzed the coupling between cadherin-based adhesions and the maintenance of collective cell coordination. We disrupted cadherin-based junctions by switching calcium levels (Fig. 3a). As expected from our previous experiments using α-catenin knockdown and Snail1-overexpressing cells, we observed that cells at confluency failed to initiate a directed rotation in low calcium medium (Fig. 3b,c,**0~21h**). Once normal calcium levels were restored, we observed a surge in coordination parameter resulting in a strong directionality followed by the establishment of well-coordinated cell rotation after 5-6 hours (Fig. 3b,c,**21~42h**). More surprisingly, when cells were switched back to low calcium conditions, the ring continued rotation even after several hours (**Movie 8**) (Fig. 3b,c,**42~51h**). A similar result was obtained by chelating calcium through the addition of EGTA during the rotation (**Extended Data Fig. 8a, b, c**). Using immunofluorescent staining, we confirmed that low calcium treatment strongly perturbed the junctional recruitment of E-cadherin without affecting focal adhesion contacts as shown by paxillin staining (**Extended Data Fig. 9**). In order to investigate the impact on single cell polarity, we analyzed the orientation of nucleus-centrosome axis with respect to the direction of rotation using calcium switch. We found a conserved polarity during collective cell rotation even after the junctions were disrupted (**Extended Data Fig. 5a,c,d**). Furthermore, we laser ablated 1-2 cells in a rotating cluster to mechanically disconnect adjacent cells. Even after the creation of a physical gap, we observed a persistent rotation of cells in their initial direction (**Movie 9**). Finally, we obtained further evidence supporting the ability of single cell to maintain a long-term polarity in the absence of E-cadherin junctions by observing single cell detachment from sub-confluent trains. These cells displayed a highly polarized persistent migration state at high speed (87 µm h^−1^), as opposed to isolated MDCK cells which did not migrate (**Extended Data Fig. 10a,b**). Altogether, these results demonstrated that the maintenance of collective cell migration was set by single cell behavior through the establishment of a sustained front-back polarity within cell clusters.

**Figure 3.**
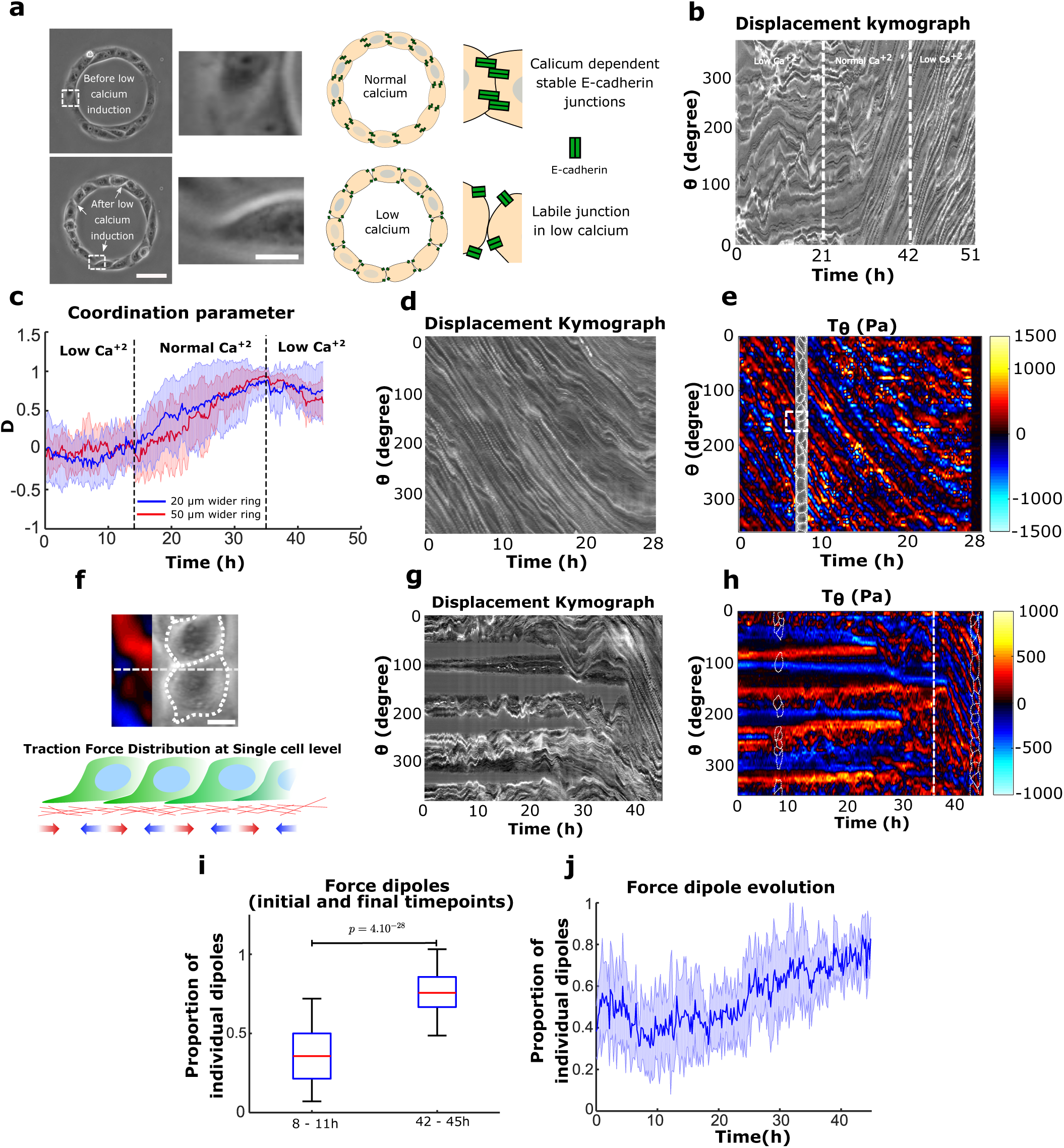
Establishment of directed collective cell migration depends on cell-cell junction strength but maintenance does not. **a**, Phase contrast images of rings under normal and low calcium conditions. The white dashed box indicates the region detailed in the expanded view (Scale bar: 10 µm); low calcium media leads to the loosening of cell-cell junctions. Schematic depicting E-cadherin distribution in normal and low calcium conditions. **b**, Representative spatio-temporal displacement kymograph of cell trains in 20 µm ring. Under low calcium cells fail to coordinate and initiate rotation (before 21 h). The normal calcium media addition (at 21 h) initiates coordinated rotations in cell train. Introducing low calcium again (at 42 h) in system maintains the coordinated rotation. **c**, Coordination parameters *D* of 20 µm (n=25, m=2) and 50 µm (n=13, m=1) wider rings indicate the similar dynamics as observed in displacement kymograph **b. d**, Kymograph of rotating cell train on soft gels for traction force measurement, traced along the circumferential midline of the ring track. **e**, Spatio-temporal distribution of tangential traction forces of rotating cell train. Straightened cell train phase contrast image overlapped with the kymograph at a given time point to map the distribution of traction forces with respect to cells in train. **f**, Expanded view, white dashed square in **e** (Scale bar: 10 µm). Schematic representing the distribution of single cell level forces. **g**, Kymograph of sub-confluent short trains (before 36 h) and confluent rotating trains (after 36 h) on soft gels for traction force measurement. **h**, Spatio-temporal distribution of tangential traction forces of multiple short trains corresponding to initial stage (before 36h) and late-stage confluent rotating ring of cells (after 36h, marked by the white dashed line). **i**, Proportion of force dipoles during initial sub-confluent uncoordinated phase (8-11h) and confluent coordinated phase of cell trains in the ring (42-45h) (n=6). **j**, Time evolution of force dipoles from initial uncoordinated state to the final rotating state of cells in ring (n=6). All scale bars unless mentioned specifically: 50 µm.

Our results challenge the classical view of collective cell migration being based on the establishment of long range propagative physical signals through intercellular contacts, as previously described for expanding cells in epithelial monolayers^3,24,37^. Therefore, we performed traction force measurements (TFM) on rotating rings and analyzed the forces applied to the substratum together with cell displacement (displacement kymographs). Interestingly, traction force kymographs exhibited distributions of force dipoles that scaled to one cell size and moved at the cellular speed (Fig. 3d,e,f). Neither the velocity nor the force fields showed a signature of long-range transmitted physical signals (**Extended Data Fig. 11a, b, c**). When PBD activity was co-registered along with TFM, we found a striking correlation between Rac1 gradient and the traction force dipoles (**Extended Data Fig. 11d,e,f,g**). We also observed that during the initial phase of uncoordinated sub-confluent cell trains, the traction forces were unevenly distributed with large forces becoming concentrated only at the free edges of the cellular clusters implicating a strong intercellular coupling, as previously described^24^ (Fig. 3g,h,i,j,**8~11h**). Forces evolved and were redistributed to establish a polarized force dipole at the single cell level after coordinated rotation was established (Fig. 3g,h,i,j,**42~45h**). We further confirmed that such force patterns were preserved in rotating rings upon calcium chelation. Again, under such conditions, we showed that mainly cell-cell junctions were perturbed with minimal impact on focal adhesions (**Extended Data Fig. 12a**). Alternatively, in α-catenin knock down cells the coordination of force dipole patterns never emerged due to lack of force coupling (**Extended Data Fig. 12b**). Interestingly, upon Arp2/3 inhibition, single cell level force dipoles were found to be diminished (**Extended Data Fig. 12c**). Overall, these findings elucidated two different phases of force distribution during collective migration: an initial phase based on large-scale forces transmission across cell clusters at sub-confluency as previously observed^3,38^, eventually leading to a phase exhibiting single cell-level force dipoles when the system reached a close state. This last phase indicates that collective cell migration can take place in the absence of long-range force transmission.

Our experimental findings led us to develop a particle-based model to validate our hypothesis regarding the respective roles of cell-cell contacts and cell polarity in driving collective rotation. We designed a minimalistic cell-based model of single cells moving on a 1D ring and driven by mechanical forces, without proliferation (see Material & Methods). In the model, cells experience viscous cell-substrate friction and cell-cell contact forces. Their migration was driven by active forces and diffusion (Fig. 4a), their polarity could adopt three states which could change in a probabilistic, force-driven way: non-migratory, migrating clockwise, or migrating anti-clockwise. Our results showed that under these minimal mechanical assumptions, cell populations spontaneously coordinated their polarity and migration and rotated over long time scales (**Movie 10a**). We analyzed this symmetry breaking process in simulations (Fig. 4b). Cells were initially non-polarized (Fig. 4b, **t=0.0h**), and after some time formed several short trains of persistently polarized cells in either direction (Fig. 4b, **t=1.0h**). Opposite trains then collided, causing the repolarization of one of the colliding trains, and finally the collective rotation of the entire ring (Fig. 4b, **t=3.3h**). This coordination process was reminiscent of our experimental observations, a result confirmed by the gradual surge of the coordination parameter to a plateau (Fig. 4c). Importantly, we found that trains with more cells have a higher persistence (Fig. 4d), confirming that forces alone can drive a length-dependent repolarization process upon frontal collisions between cell trains, as seen in our experiments and in previously reported study^39^.

**Figure 4.**
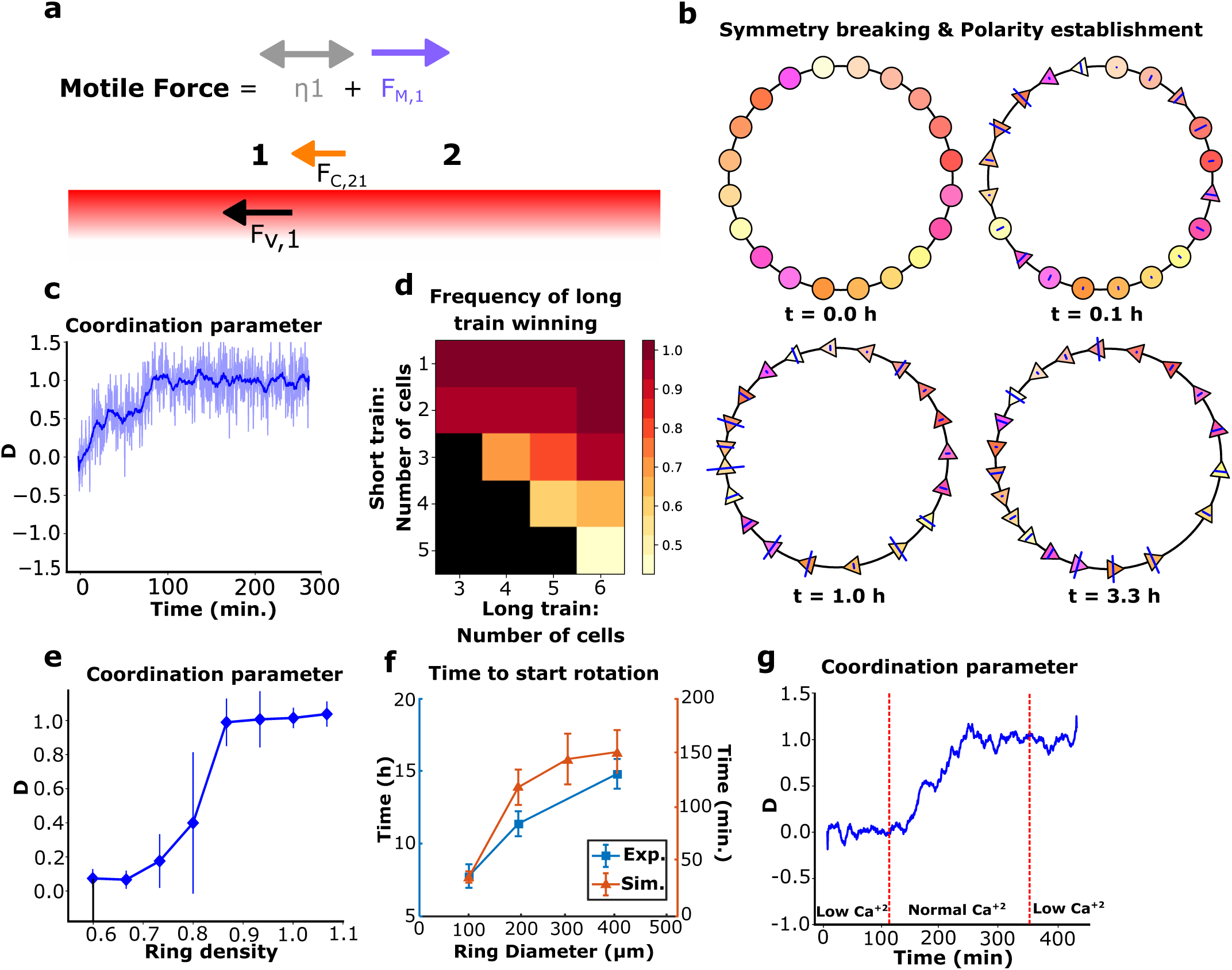
Mechanics-based simulations of epithelial cell migration on a ring. **a**, Schematic of force balance in epithelial cell train. The forces are F_V_ (viscous), F_C_ (contact), F_M_ (directed motile force) and the diffusion ŋ. **b**, Coordination emergence in a simulation. Circles: centers of non-polarized cells. Triangles: centers of polarized cells, pointing in their polarity direction (cell boundaries are not represented). Blue lines indicate the intensity of contact forces on a cell. **c**, Simulations reveal that persistent coordination emerges over time (blue pale line: coordination parameter for n=1 simulation, smoothed over time in the blue thick line). **d**, Colormap of the probability for the longer train to “win” a collision versus the number of cells in both trains. Black square: no data. n=20. **e**, Coordination parameter in simulations after 1000 timesteps (~400 minutes, averaged over last 8 minutes) versus cell density 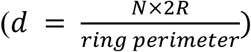 demonstrates that confluent density is necessary to achieve coordination. n=20 per point. **f**, Coordination time in experiments (n=20) and simulations (n=25) with varying ring sizes. **g**, Simulations varying the contact stiffness parameter over time to modulate cell-cell interactions (blue pale line: coordination parameter for n=1 simulation, smoothed over time in the blue thick line). *ring diameter is 200 µm unless stated otherwise.

Furthermore, as observed in experiments, coordination required cells to be under confluent density: when intercellular interactions were limited by a low global density, collective rotation failed to emerge (Fig. 4e). Again, consistent with the experimental data, persistent rotation emerged independently of the annular ring size, confirming that single-cell behaviors allow the long-range transmission of information over the entire system. Coordination times in both model and experiments (defined in Material & Methods) increased with the ring diameter (Fig. 4f). To compare the dynamics of cell coordination in our model with experimental data, the spatial velocity correlation function on the ring was calculated (**Extended Data Fig. 13a**). We observed striking similarities between both patterns: periodic patterns larger than one cell progressively appear, suggesting the gathering of cells in several polarized trains, and disappear with time as cells migrate collectively. To study the importance of cell-cell junction-based interactions as experimentally determined through switching calcium levels, we decreased the ability of cells to repel each other upon contact (low numerical contact stiffness), facilitating as a result their movement past each other (**Movie 10b, top**) as compared to the case of cell collision based repolarization under normal calcium (normal numerical contact stiffness) (**Movie 10b, bottom**). Remarkably, this alone was sufficient to recapture the effect of low Ca^2+^ levels: the initiation of coordination was made impossible, while collective movement was not impaired in already coordinated cell populations (Fig. 4g). Finally, simulations exhibited a transition in the migratory modes: cell-substrate friction is balanced by intercellular interactions during train formation and merging, and by motile forces when cells are fully coordinated (**Extended Data Fig. 13b**), which is reminiscent of the transition in force dipoles we observed experimentally. Taken together, these results indicate that our model successfully capture the overall dynamics of coordination emergence, highlighting the importance of single cell polarity and intercellular interactions in the resulting persistent rotation.

Collective cell migration has been described so far through the indispensable role of long-range interactions, the establishment of firm cell-cell junctions and supracellular coordination^3,33,40^. However, even shown in *in vitro* and *in vivo* studies^6,9^, the role of single cell behavior in multicellular systems has been underestimated. Here, we show that the maintenance of collective cell migration in monolayers lacking a free front edge relies on the emergence of single cell mechanical entities that coordinate their front-back polarity with the extension of lamellipodial-based structure underneath the front cell. Our findings reveal that this robust process relies on a long-term memory of the cellular polarized state which, once formed, becomes independent of cell-cell junctions. This driving mode of collective cell movements can recapitulate several situations observed *in vivo* during morphogenesis^5,6,41,42^, shed new light in the interpretation of previous studies by taking into account the local behavior of single cells and thus provide new mechanism to interpret directed cell migration during development, wound healing, and collective cancer cell invasion.

## Acknowledgements

The authors thank Delphine Delacour, Jacques Prost, Masahiro Sokabe, Yusuke Toyama, Raphael Voituriez, and group members from MBI and IJM for helpful discussions and critical reading of the manuscript. We would like to thank MBI science communication team member Steven Wolf and Andrew Wong for the proof reading of manuscript. We also thank the MBI Microfabrication Core (Gianluca Grenci and Mohammed Asraf) and MBI Microscopy Core (Felix Margadant) for continuous support and would like to extend our thanks to Ong Hui Ting for her valuable suggestions in image processing and image analysis. We also acknowledge the ImagoSeine core facility of the Institut Jacques Monod, and members of IBiSA and France-BioImaging (ANR-10-INBS-04) infrastructures. The authors are grateful to W. J. Nelson and F. Martin-Belmonte for their generous gift of MDCK cell lines.

## Author contributions

B.L., R.M.M and C.T.L supervised the project; S.J. and B.L. conceived the study and designed the experiments. S.J., G.H.N.S.N., V.Cel., S.D.B., M.H. and T.C. performed experiments; S.J.,T.C., J.A, G.H.N.S.N., did experimental data analysis; RMM and P.M. contributed new reagents, modeling and computational tools; V.C. and A.J.K. implemented the numerical method and did the *in-silico* simulations. B.L., S.J. and V.C wrote the manuscript. All authors critically proofread the manuscript.

## Data and materials availability

Analysis tools are available for purposes of reproducing or extending the analysis.

