## Supplementary material for "Emergence of single cell mechanical behavior and polarity within epithelial monolayers drives collective cell migration": Methods and Supplementary figures

**This PDF file includes:**

Materials and Methods
Supplementary Text
Extended Data Figures 1-14
Movies Caption 1-10

**Other Supplementary Materials for this manuscript include the following:**

Movies 1-10

#### Material and Methods

##### Cell Culture, Microcontact printing and Cell Seeding

MDCK-WT, MDCK-Histone-GFP, MDCK-Ecadherin-GFP and MDCK-PBD-YFP cells were maintained in DMEM (Dulbecco's Modified Eagle Medium) (Invitrogen) supplemented with 10% FBS (Fetal Bovine Serum) (Invitrogen) 1% Penicillin-Streptomycin antibiotic (Invitrogen). All cells were maintained at 37°C and 5% $CO_2$. Cells were passaged after reaching 70% confluency. The MDCK opto-Rac1 stable cell lines was obtained by lentiviral infection: lentiviral particles were produced by transfecting the pHR-TIAM-CRY2-mCherry or pLVX-CIBN-GFP-CAAX plasmids along with the vectors encoding packaging proteins (pMD2.G and psPax2) into HEK-293T cells. Viral supernatants were collected 2 days after transfection and MDCK cells were transduced at a MOI of 2. Clones with appropriate expression levels of TIAM-CRY2-mCherry and CIBN-GFP-CAAX were then selected.

Roughly 0.7 x $10^6$ cells were seeded on fibronectin (Sigma) patterns[25] placed in a 35mm PDMS (Sylgard 184, Dow Corning) spin coated glass dish. Cells attached on fibronectin ring patterns within 45 min. to 1 hr. Cells were then washed once with DPBS (Dulbecco's phosphate-buffered saline) (Gibco) and once with DMEM media to prepare the sample for final imaging.

##### Microscopy

Long-term-time-lapse imaging to observe large scale epithelia dynamics was done using Nikon Biostation IMQ. We used 10X,20X objectives with NA 0.5 for imaging.
To observe the live spatio-temporal dynamics for the protein of interest, Nikon A1R confocal, Nikon CSU W1 spinning disk and Zeiss LSM 780 confocal were used. Most confocal images were taken with 60X/1.40 oil-immersion objective. Same set of confocal microscopes were used for acquiring immunofluorescence images.

##### Drug treatment

In most experiments the cell cycle was blocked using Mitomycin-c (Sigma Aldrich) added to the culture at 10 µg/ml for 1 hour, followed by washing twice with DPBS and twice with DMEM. The Arp 2/3 inhibitor, CK-666 (Sigma Aldrich) was used at 100 µM. The myosin II ATPase inhibitor Blebbistatin (Sigma Aldrich, B0560-1MG) was used at 80 µM. The Rac1 inhibitor Drug Z62954982 (Merck Millipore) was used at 100 µM. Primarily, to block cadherin adhesion, the entire normal media was replaced with low calcium media (~20 µM calcium) after 2 consecutive wash from low calcium media to remove any left-out calcium residue because of the prior existing normal media. Alternatively, EGTA (Sigma Aldrich) was added to the normal medium at a final concentration of 2 mM on our regular experiments with stiff substrate (spin-coated glass PDMS dishes) to partially chelate the calcium in the media. For the low calcium experiments done on soft substrates, we used EGTA concentration of 1.8 mM.

##### Preparation of low calcium media

Low calcium media was prepared by adding 1% FBS, 1% Penicillin-Streptomycin, 1% Sodium Pyruvate (Invitrogen) and 1% GlutaMAX (Gibco) to DMEM without calcium (Gibco). The final calcium concentration of the complete DMEM was approximately 20 µM.

**Laser ablation of cells**

We used a custom-built laser system integrated with a Nikon A1R MP laser scanning confocal microscope to ablate the cells in the cell monolayer through a Nikon apo 60X/1.40 oil-immersion objective. The laser used in the ablation system was in UV range with a pulse duration of 300 ps and a repetition rate of 1 kHz. The Power-Chip details: PNV-0150-100, Teem Photonics. We used a laser power of ~ 100 nW to 200 nW and the exposure time ranging from 0.5 seconds to 2 seconds to target cell nucleus in MDCK histone GFP stable cells to instantly ablate or blast the cell to make the epithelia discontinuous. Followed by the ablation, the images were acquired on the same microscope at 60X magnification. MDCK histone GFP stable cells were generously gifted from Sham Leilah Tlili (Mechanobiology Institute, Singapore)[43].

**Displacement and Velocity Kymographs:** To obtain displacement kymographs, a centerline across the ring width is traced along its circular perimeter. Standard ImageJ RESLICE plugin is then used for the entire experimental movie to obtain a spatiotemporal displacement kymograph. To obtain the velocity field, we used an open source PIV (particle image velocimetry) MATLAB code named MatPIV[44]. To generate respective kymographs for obtaining a spatio-temporal distribution of velocities, the ring was spatially divided into 18 segments of 20° for each timepoint. The velocities were averaged in each segment to obtain a spatial profile. Similar steps were performed for every timepoint to obtain a heatmap representing a spatio-temporal distribution of the required dataset. Each segment of 20° is approximately equal to 31 µm.

**Traction Force Microscopy (TFM) and traction force kymograph**

The substrate for Traction Force Microscopy was prepared using soft silicone gel mixed in the ratio 1:1 (CyA:CyB) (Dow Corning). The method has been described in previous studies[45,46]. Briefly, a thin layer of the gel was spread on a glass bottom dish and then cured at 80°C for 2 hours. Cured gel was silanized using 5% solution of (3-aminopropyl) triethoxysilane (APTES, Sigma) in pure ethanol. This gel was later incubated for 5 minutes with 100 nm carboxylated fluorescent beads (Invitrogen) suspended in deionized (DI) water. Subsequently, the substrate was dried, and micro-contact printed with fibronectin using a thin water soluble polyvinyl alcohol (PVA) membrane which allows the transfer of fibronectin stamps on soft gel[25,47]. The PVA membrane was later dissolved and the non-contact printed areas were blocked using 2% Pluronics (Sigma) solution. The substrate was then washed and seeded with cells. Cells were allowed to grow until the micro-contact printed area reached confluency.

The samples were imaged using a Biostation IM-Q (Nikon) time lapse imaging microscope at 20X magnification for several hours. Images were acquired using the phase contrast and fluorescent channels to record images of the cell along with bead displacement.

For analysis, the imaging drifts were corrected in ImageJ (NIH) using the Image Stabilizer plugin[48]. To analyze the displacement field of beads, we used an open source PIV (particle image velocimetry) MATLAB code named MatPIV[44]. To reconstruct the traction force field from the obtained displacement field, an open source Fourier transform traction cytometry (FTTC) plugin was used in ImageJ[49]. To construct the final heat map of the tangential traction forces, the ring was divided into 72 equal sectors of 5° each and the values were averaged over the entire ring space for every 5° segment. The same was done for the entire time frame to plot spatio-temporal plots. Each segment of 5° is approximately equal to 7.85 µm.

**Immunofluorescence and cell transfection**

Cells were fixed with 4% formaldehyde in PBS at 37°C followed by blocking for 1 hour in a Saponin and BSA mixture at room temperature. In the case of centrosome immunofluorescence, methanol was used for fixation at -20°c. Cells were then rinsed with PBS 3 times. Subsequently, cells were incubated with primary antibodies for 1 hour followed by a PBS wash. Secondary antibodies were further incubated for 1 hr. After washing thoroughly with PBS 3 times, cells were mounted with coverslip and Vectashield mounting media (Vector laboratories) for imaging. Primary and secondary antibodies used for immunofluorescence are listed below:

| Primary Antibody | Company | Dilution |
|---|---|---|
| Rabbit polyclonal anti-paxillin – ab32084 | Abcam | 1:100 |
| Rat monoclonal anti-E-cadherin – U3254 | Sigma | 1:100 |
| Mouse monoclonal anti-ϒ -Tubulin – T6557 | Sigma | 1:200 |
| Rat monoclonal anti-Ecadherin (clone DECMA1) – MABT26 | Sigma | 1:200 |

| Secondary Antibody | Company | Dilution |
|---|---|---|
| Alexa 488 conjugated donkey anti-rat IgG | Invitrogen | 1:100 |
| Alexa 568 conjugated goat anti-rabbit IgG | Invitrogen | 1:100 |
| Alexa 568 conjugated goat anti-mouse IgG | Life Technologies Ltd. | 1:200 |
| Nuclei staining – Hoechst reagent | Invitrogen | 1:1000 |

**Cell persistency calculation**

Persistence is defined as the probability of the cell's velocity vector to remain in the same direction (either clockwise or anti-clockwise on the ring) between consecutive imaging frames (10 minutes per frame), calculated over the available time-lapse movie when a single detached cell is observed, according to the following formula:

Persistence = $\sum_{0}^{n=frame} \frac{1\ if\ velocity\ vector\ is\ same\ direction\ as\ previous\ frame, 0\ otherwise}{n}$.

Hence, persistence=1 when cells do not change direction, persistence=0.5 when cells change direction randomly, persistence=0 when cells engage in oscillatory motion with the sampling frequency.

**Error bar and statistical testing**

All error bars indicated in plots are the Standard deviations and the statistical testing are done using unpaired two tailed t test unless it is mentioned in some specific cases where unpaired one tailed test was performed. In figures, **n** represents total number of points considered (total number of rings, or cells respectively based on experimental requirement) and **m** represent the number of experiments. Wherever, **m** is not mentioned, it indicates one experiment set or a representative image.

**Measurement of coordination time for different ring size**

Cells were seeded under low calcium conditions on different sized ring micro-patterns to ensure that all rings reached nearly a full confluency after initial cell seeding. The low calcium media (~20 µM) ensured that cells were not able to initiate coordination dynamics in absence of calcium dependent adherens junction formation. Cells were imaged in low calcium media for few hours before introducing the normal calcium media (2 mM). This protocol ensured that we present the same switching point to initiate coordination in all the different sized rings at confluent condition. We started out time recording from the point of introducing normal calcium media until the time at which each single cell in the ring began to rotate in one direction. We stopped and recorded our time measurement at this point. This time recorded was referred as the coordination time for different ring size diameter respectively. These experiments were performed with MDCK histone GFP stable cells to facilitate the ease of cell tracking.

**Transfections:**

For simultaneous visualization of PBD and AHPH dynamics, YFP-PBD MDCK cells[50] were transiently transfected with a plasmid driving the expression of mcherry AHPH (generous gift from Alpha Yap, The University of Queensland, Australia)[32,51], Two million YFP-PBD MDCK cells were electroporated (Neon Transfection System Invitrogen) with 3-5 ug of plasmid encoding AHPH-mcherry in one pulse of 20 ms at 1650 V. After transfection cells were seeded directly on the ring patterns and live imaging was performed after 24 hours, with a Zeiss LSM 780 confocal microscope equipped with a 63 X oil immersion objective at a resolution of 0.8 µm z-stacks in a range of 6-7 µm, time-lapse was done for 8 hours at a rate of 1 frame every 10min.
The pHR-TIAM-CRY2-mCherry, pLVX-CIBN-GFP-CAAX and PBD-iRFP plasmids used in optogenetics experiments were a kind gift of Mathieu Coppey[52].

**Vertical confinement of lamellipodia:**

To check the hypothesis of confinement mediated increase in lamellipodia activity. We designed a strategy based on our serendipitous observations during cell migration experiments earlier, where we confine cells by placing a PDMS block and latter releasing this physical confinement that allows cell to migrate. Our observations were that cells even though they are strictly confined and do not enter the PDMS block, but at times in few locations the lamellipodia would penetrate under the PDMS block but not the whole cell. We took advantage of this observation and devised a strategy to check our hypothesis. Here we placed a PDMS block on the glass bottom dishes coated with fibronectin for 1 hour with 50 µg/mL human plasma fibronectin (Merck Millipore) diluted in PBS and then washed three times with PBS. Then GFP-Actin cells were seeded on these dishes with confinement, were allowed to spread and live imaging was done to capture the events where lamellipodia has penetrated under the PDMS block (**Extended Data Fig. 4**). Live imaging was done with a Zeiss LSM 780 confocal microscope equipped with a 63 X oil immersion objective at a resolution of 0.15 µm z-stacks in a range of 2-2.5 µm, time-lapse was done for 5 min at a rate of 1 frame every 30 seconds. **Analysis :** For intensity profiles, a line was drawn on the ROI and the line scan was done using the plot profile plugin in ImageJ, the values obtained were then plotted. Graphs and statistical tests were done using GraphPad prism software.

**Optogenetics experiments**

Trains of opto-Rac1 cells were initially left to rotate on ring patterns. In order to perturb the front-rear Rac1-based polarity, we illuminated a region of interest corresponding to a quarter of the circumference of these rings with blue light (405 nm). Uniform activation of Rac1 in the cells in these regions resulted in a loss of cell polarity[52] (**Extended Data Fig. 7a,b**). The loss of Rac1 gradients could be verified by the flattening of PBD-iRFP signal (**Extended Data Fig. 7a,b**).

The microscopy was performed using Zeiss Axio Observer Z1 mounted with spinning disk CSU-X1 (Yokogawa) unit. The photoactivation was performed using FRAP unit with 405 nm laser.

**Analysis of ring closure**

To determine what criterion predicted best the ring's direction of rotation, we counted the number of rings in which the condition related to the given criterion was satisfied, such as for the criterion "train length" the condition was "the ring finally rotates in the same direction as the longer train". The criteria and conditions were the following:

**N.** 3 frames before (i.e 30 minutes before) ring closure, the train was divided in two, oppositely moving, parts. The cells in each part were manually counted using the transmission images and the condition tested was that the ring finally rotates in the same direction as the part with the larger number of cells.

**<v>.** The train was divided in two parts as before. The average (absolute value of the) speed <v> was computed in the two parts. The condition tested was that the ring finally rotates in the same direction as the part with larger <v>.

**Vfront.** The velocity was measured in a sector of 12.5° (44 µm) next to both train ends. The condition tested was that the ring finally rotates in the same direction as the faster end.

**Tfront.** The traction forces were measured as described, and their magnitude was averaged on a zone next to the end. Contrary to the velocity, this zone was manually defined on traction force kymographs to engulf the whole traction patch of the end. The condition tested was that the ring finally rotates in the same direction as the end with the larger traction magnitude.

**Analysis of dipoles**

The rings were first segmented using binarized transmission images to identify coherent trains. The number of dipoles at a given time was determined as the number of 0-crossings of the T(theta) curve along the ring at this time, excluding the areas that were not covered with cells. Then, the proportion of internal dipoles was obtained as follows: (i) the number of internal junctions $N_{ij}$ was defined as $N_{ij} = N_c - N_{holes}$, where Nc is the number of cells in the ring and $N_{holes}$ the number of holes between trains. (ii) The number of internal dipoles was defined as $N_{id} = N_d - N_{holes}$, where Nd is the total number of dipoles. Finally, the proportion of internal dipoles is $P_{id}=N_{id}/N_{ij}$, so that it falls to 0 if all trains contract as units, and it goes to 1 if all cells contract individually even when they have cell-cell junctions.

**Spatial correlation of traction forces and PBD signal**

To quantify the spatial relationship between the traction force pattern and the cell polarity, we computed the normalized spatial cross-correlation coefficient defined as:

$$C_{f,g}(\Delta\theta) = \frac{\langle \tilde{f}(\theta+\Delta\theta).\tilde{g}(\theta)\rangle_\theta}{\left(\langle \tilde{f}(\theta)^2\rangle_\theta.\langle \tilde{g}(\theta)^2\rangle_\theta\right)^{\frac{1}{2}}}$$

Where the angle brackets denote average, $\tilde{f} = f - \langle f \rangle_\theta$ and $\tilde{g} = g - \langle g \rangle_\theta$, $f$ is the orthoradial traction force, $g$ is the PBD signal, and $\theta$ is the position along the ring. All the rings were first aligned so that they all rotate in the same direction. Then this function was computed for each time frame and averaged over time for each ring. Finally, the plot shows the average over the 6 rings and the error bars denote the standard deviation.
To check that the peak appearing close to $\Delta\theta = 0$ does not stem accidentally from the signals' properties, we shuffled one or both signals as follows. We picked a random phase-shift $\theta_{shift}(t)$ (all independent for various times, rings and for $f$and $g$), and transformed the signal as $f_{shifted}(\theta, t) = f(\theta + \theta_{shift}(t), t)$. Then we proceeded as before, replacing $f$, $g$, or both $f$ and $g$ by their shifted version. That way we conserved the original signals' structure but removed their potential correlations. Indeed, the resulting curves match the plot in **Extended Data Fig. 11g** for $\theta < 90°$ or $\theta > 0°$ showing that the peak for slightly negative $\theta$ denotes a significant correlation between the two signals.

**Analysis of nucleus-centrosome axis orientation**
After the confluency when rotation begins, the rings were allowed to rotate for ~24 hours. The sample was imaged for 2 hours for phase images with the help of Biostation IMQ (Nikon). Taking any one ring present at corner of sample as our reference, we noted the direction of rotation of other rings. The samples were later fixed, and immunofluorescence was performed on them using the ϒ-Tubulin antibody (listed in the antibody table) and nucleus was already labelled in GFP since we used MDCK-Histone-GFP cell line for this purpose. Further the alignment of centrosome and nucleus was recorded for every cell of the rings with respect to the rotation direction of the particular ring respectively. If the centrosome was placed before the nucleus in a way such that if the migration direction arrow tail is represented by the nucleus and the arrowhead is represented by the centrosomes, then this configuration is taken as CN mode or aligned mode (Centrosome placed before Nucleus). In other case, if the arrow of migration direction tail represents Centrosome and the arrowhead represents the Nucleus then this is NC configuration or not-aligned mode (Nucleus placed before centrosome).In all the other cases, where there are multiple centrosome present because of a dividing state of a cell or if the Nucleus-centrosome are aligned in the perpendicular direction of the migration direction arrows, then this is taken as undetermined condition.

For the cases where rings are not rotating in any definite direction, for example in the case of under confluent rings and the low calcium condition (before symmetry breaking or before the rotation). We chose a reference direction if the ring rotated in clockwise direction and hence the Nucleus-centrosome alignment was computed in the same way as explained above. We indeed also verified the other case by taking reference direction as anti-clockwise and we didn't find any positive correlation same as the clockwise case. Further robust characterization we performed in order to validate our hypothesis for the analysis of non-rotating rings are not included in the results.

**Supplementary Text**

**Numerical model**
A force-based model of active cells was implemented to study emergence of collective rotation on rings of cells. A one-dimensional framework was used, modelling the ring as a segment with periodic boundaries associated to an *x*-axis.

Our model is entirely driven by explicit mechanical processes, with the single cell as the building block. Cells are segments of original length $2R_0 = 40\mu$m, located using the position of their center on the *x*-axis. The use of a force-balance method allows the explicit tracking of individual cells. Cells are subject to three types of surface forces: viscous cell-substrate forces proportional to their velocity; elastic contact forces representing volume exclusion; and active forces.

Active forces are implemented as the sum of a noise $\boldsymbol{\eta}$, representing autonomous cell diffusion, and an effective persistent migration force $\boldsymbol{f_M}$. $\boldsymbol{\eta}$ is a 1D Gaussian noise of mean 0 and standard deviation σ. $\boldsymbol{f_M}$ strongly relates to cell polarization: cells in our model can be found in one of three polarity states (*S, CW* and *CCW*) representing their possible dynamical behaviors. The migration force $f_M$ is equal to 0 for a non-persistently polarized cell, referred to as *static (S)* and exhibiting Brownian motion at the considered time scale; it is equal to $\pm F_M$, where $F_M$ is a constant, for a persistently polarized cell in either direction, referred to as *clockwise (CW)*- or *counter clockwise (CCW)*-polarized.

The contact force caused by cell *i* on another cell is:

$$\boldsymbol{f_{c,i}} = F_C\left(1-\frac{d_i}{2R}\right)1_{\{2R-d_i\}}\boldsymbol{u_i}$$

where $F_C$ is the cell stiffness, *R* the cell radius, $d_i$ is the distance between both cells and $\boldsymbol{u_i}$ is the unit vector pointing from cell *i* to the other cell. $1_{\{x\}}$ is the Heaviside step function, equal to 1 if $x \geq 0$ and to 0 otherwise. $F_c$ is an effective parameter representing the extent to which cells will react, reorient and change their behavior upon intercellular contact: it encompasses volume exclusion due to mechanical stiffness, but also biochemical parameters such as intercellular adhesion.

Thus, the total balance of forces on a cell migrating on the ring with velocity $\boldsymbol{v}$ is:

$$\mu\boldsymbol{v} = \left(f_{C,i} - f_{C,j}\right)\boldsymbol{u} + \boldsymbol{f_M} + \boldsymbol{\eta}$$

where $\mu$ is the viscous friction coefficient, cells *i* and *j* are the nearest left-side and right-side cell neighbors respectively, and $\boldsymbol{u}$ is the unit vector pointing towards the positive *x*-axis.

[Desai et. al,2013, RSI] showed that CIL-associated cell repolarizations occur probabilistically, an observation we adopted in our model. In our model, the probability for a cell polarity to change is mechanically driven: it depends on the forces on this cell and is controlled by a force threshold $F_0$ [28]. Single isolated cells in experiments never get persistently polarized, indicating that $\boldsymbol{\eta}$ does not impact polarity changes. Moreover, repolarization requires intercellular contacts lasting a few minutes, indicating that an average force during a contact time $\tau_{int}$ is more likely to drive repolarization than a high, punctual force. Therefore, the probability for a cell to become persistently polarized at a given time *t* increases with $\frac{1}{F_0}\left|\int_{t-\tau_{int}}^{t} f(u)du\right| \equiv \frac{1}{F_0}\int f(t)$, where $f$ is the projection of the sum of $\boldsymbol{f_M}$ and of contact forces on the cell. It is worth noting that persistent polarization seems to be irreversible: return to a *static* state is impossible.

For a cell in the polarity state *X* subjected to $\int f$, for a given time step *dt*, the probability distributions associated to polarity changes are given by:

$$\mathbb{P}(X \to CCW|\textstyle\int f, dt) = 1 - \left(\dfrac{1}{1+e^{k_s\left(\frac{\int f}{F_0}-1\right)}}\right)^{\frac{dt}{\tau_{int}}} \underbrace{+\,\mathbb{P}_{remain}(CCW|\textstyle\int f, dt)}_{\text{if } X=CCW}$$

$$\mathbb{P}(X \to CW | \int f, dt) = 1 - \left( \frac{1}{1 + e^{-k_s\left(\frac{\int f}{F_0}+1\right)}} \right)^{\frac{dt}{\tau_{int}}} \underbrace{+ \mathbb{P}_{remain}(CW | \int f, dt)}_{\text{if } X=CW}$$

$$\mathbb{P}(X \to S \mid \int f, dt) = \begin{cases} \mathbb{P}_{remain}(S | \int f, dt) \text{ if } X = S \\ 0 \text{ otherwise} \end{cases}$$

Where $\mathbb{P}_{remain}(X | \int f, dt) \approx \left( \frac{1}{1+e^{k_s\left(\frac{|\int f|}{F_0}-1\right)}} \right)^{\frac{dt}{\tau_{int}}}$ is a sigmoid function of steepness $k_s$.

Biophysical parameters were calibrated using experimental data. The persistent migration parameter $\frac{F_M}{\mu}$ was set using the average cell velocity during collective migration.

The cell diffusion parameter $\sigma$ was adjusted using previous single cell literature. In 1D, the random motility coefficient of cells as defined in[53] is analogous to the cell diffusivity $\frac{1}{2}\left(\frac{\sigma}{\mu}\right)^2$ and equals $S^2P$ where $S$ is the root mean-squared cell velocity and $P$ is the correlation time of cell direction. Literature for single MDCK cells provided us with orders of magnitude for S and P[54,55] and thus for $\sigma$ relative to $\mu$.

The parameters $k_s$ and $F_0$ characterizing the probabilistic polarity changes were chosen so that the polarization time for a single cell hit by a 5 cell-train matches experimental values, and the probability for an isolated static cell to become persistently polarized is negligible. The cell stiffness $\frac{F_C}{\mu}$ was estimated by measuring cell compressibility in experiments of polarized cell trains migrating between obstacles (experimentally, obstacles were the borders of a non-continuous micro-pattern fibronectin ring) (**Extended Data Fig. 14a**). During the collective migration of a cell train towards the obstacle, the cell at the rear end of the train was seen to migrate unconstrained, with a radius $R$. When the front cell was stopped by the obstacle, the rear end cell continued migrating and got gradually compressed against the immobilized train (**Extended Data Fig. 14b**). Its velocity decreased until the cell stopped, with an equilibrium radius $R_e$ (**Extended Data Fig. 14c, d**). Let us analyze the behavior of the rear end cell in terms of our model. Using force balance, we can deduce that $RF_M = (R - R_e)F_C$. By measuring experimental cell compression, we thus get an estimate of $F_C$ relatively to $F_M$.

The model was implemented using Euler's method with a 24 second-timestep. Simulations were run up to 1 000 steps to reach a steady state.

**Table of model parameters:**

| Parameter | Definition | Value |
|---|---|---|
| L | Ring perimeter | 600 (μm) |
| R | Cell radius | 20 (μm) |
| $\mu$ | Viscous friction | No data |
| $F_M$ | Persistent motile force | 8.3 $10^{-9}$ $\mu$ (Pa) |

| | | |
|---|---|---|
| σ | Cell diffusion parameter | 4.5 $10^{-8}$ $\mu$ (Pa) |
| $F_C$ | Cell stiffness | 4.6 $10^{-8}$ $\mu$ (Pa) |
| $\tau_{int}$ | Contact time | 60 (s) |
| $k_s$ | Sigmoid steepness | ~12 |
| $F_0$ | Force threshold | 2.0 $10^{-9}$ $\mu$ (Pa) |

Coordination Time

The coordination time was defined as the duration between the last time of no preferential direction on the ring ($D$ = 0) and the time when the coordination parameter reaches a final, lasting plateau at ±1, indicating complete coordination of the cells.

Correlation function

The spatial velocity correlation function was calculated for both experiments and simulations using the formula below:

$$C(x',t) = \frac{< u(x+x',t) \times u(x,t) >_x}{\sqrt{< u(x+x',t)^2 >_x \times < u(x,t)^2 >_x}}$$

where $x$ and $x'$ are curvilinear abscissas along the ring, $u$ is the angular velocity (positive in the counter-clockwise direction) and $t$ is time.

### Extended Data Figure 1

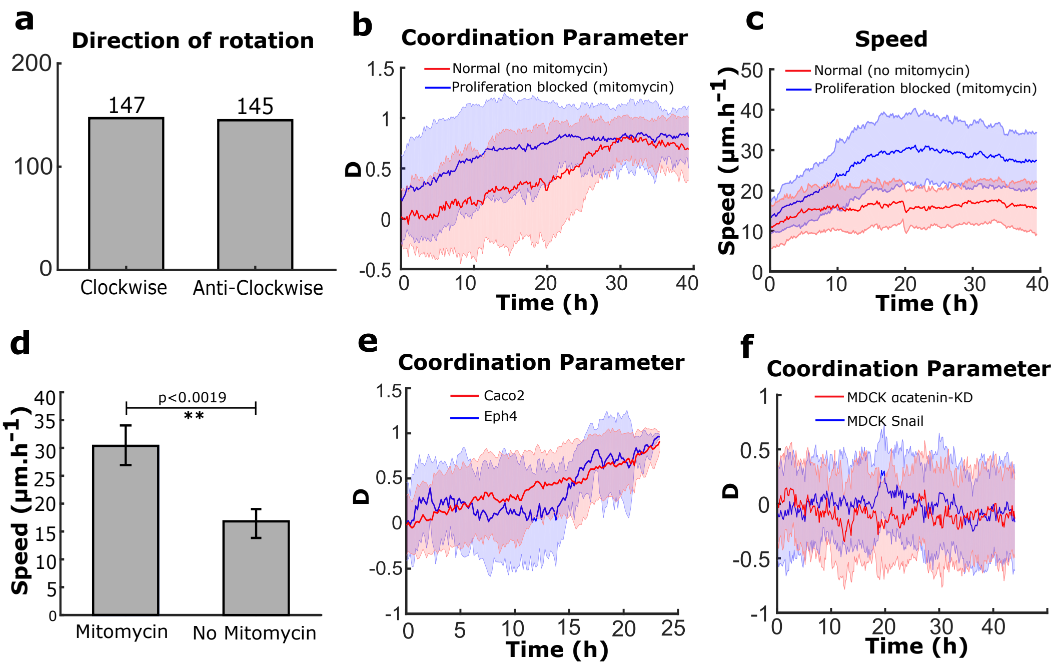

### Extended Data Figure 2

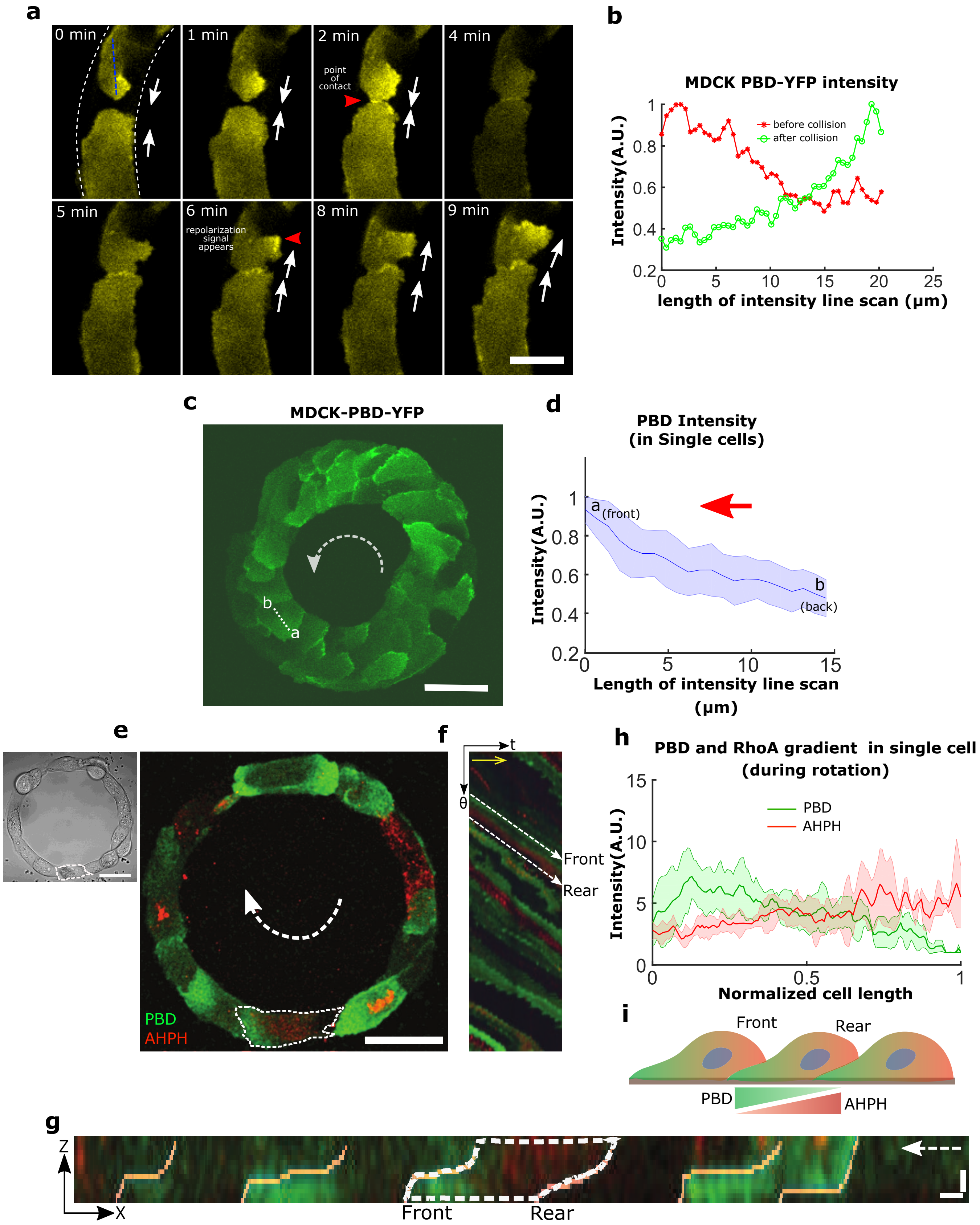

### Extended Data Figure 3

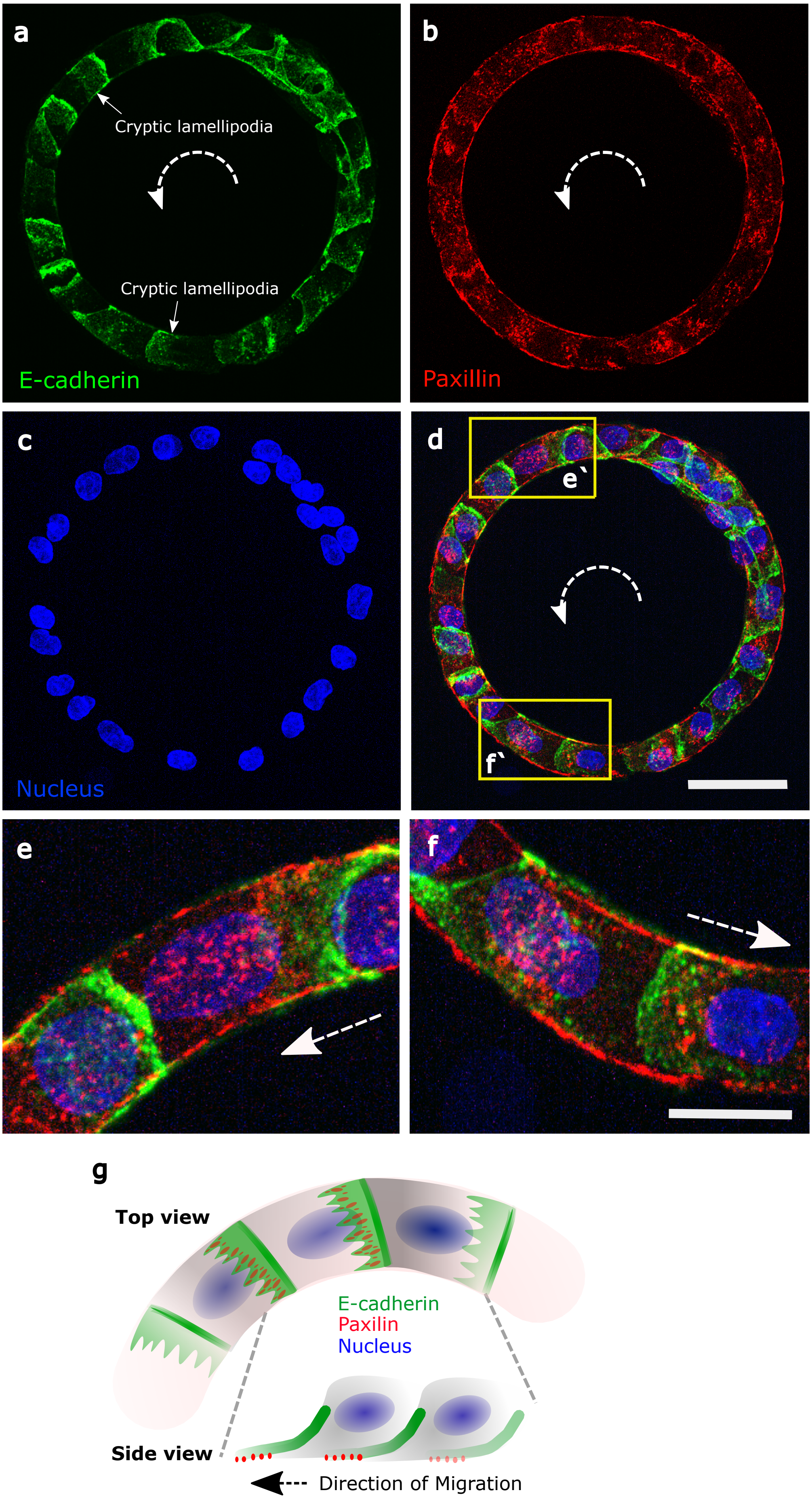

### Extended Data Figure 4

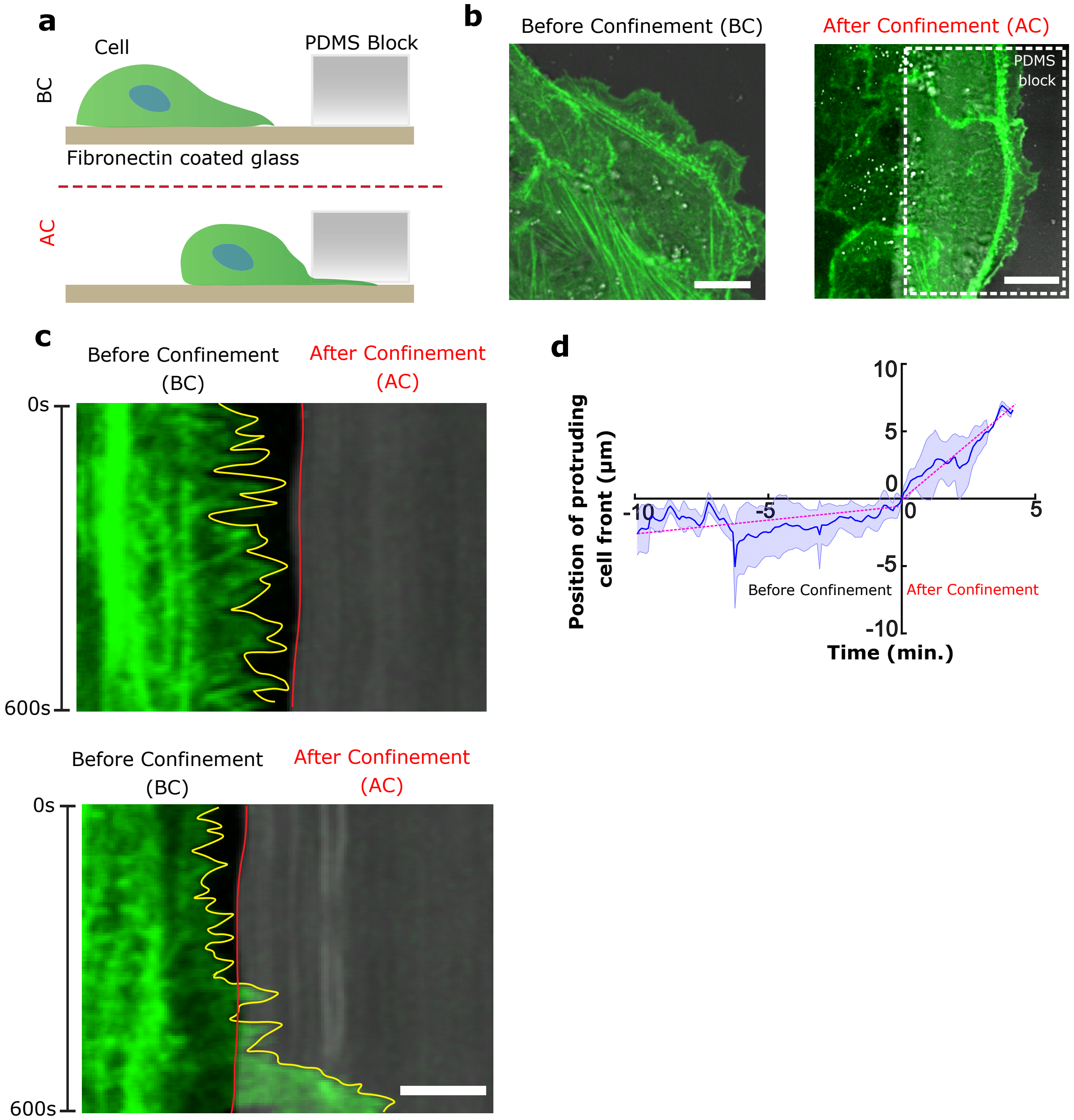

### Extended Data Figure 5

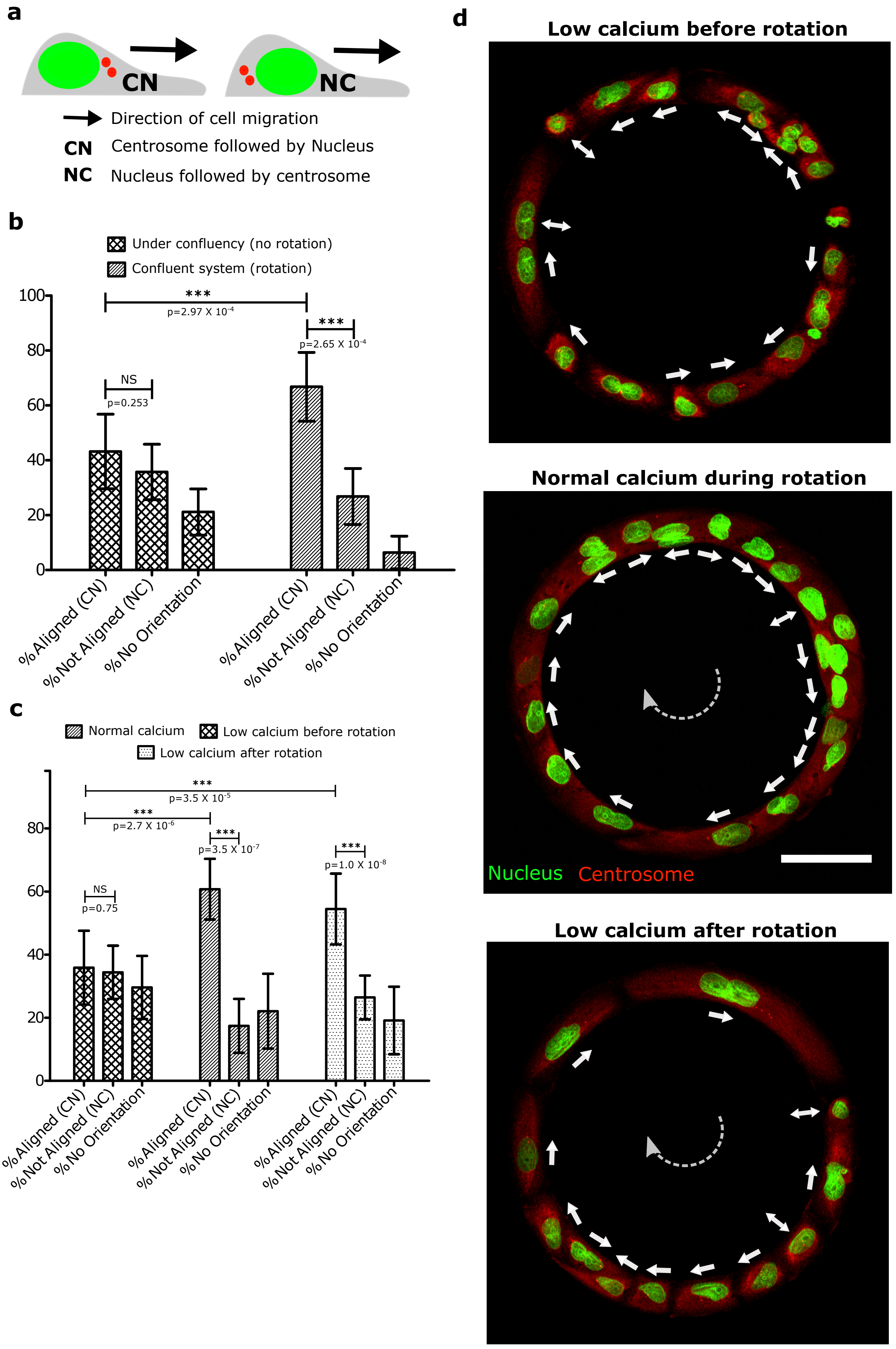

### Extended Data Figure 6

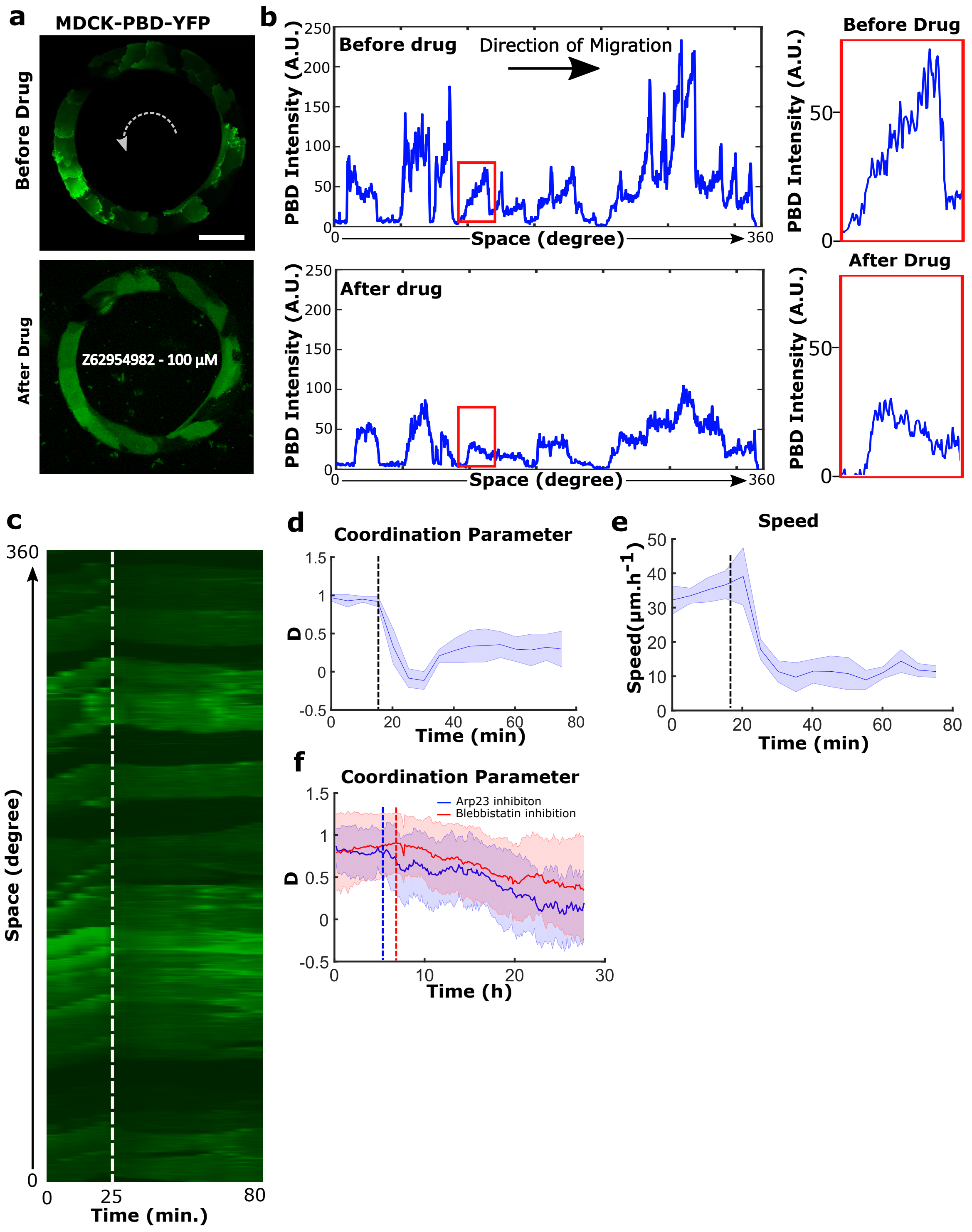

### Extended Data Figure 7

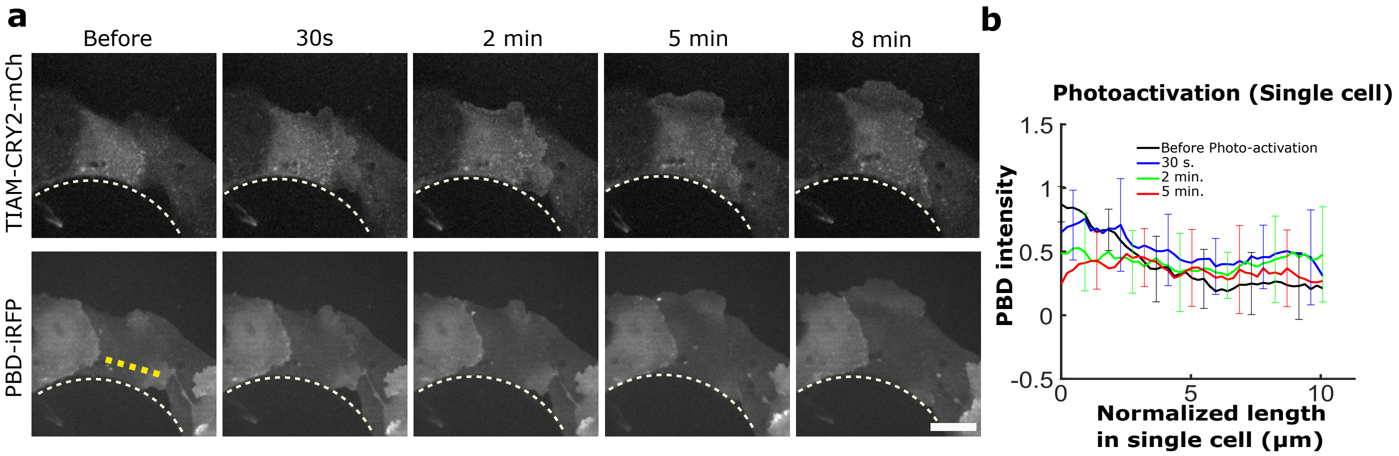

### Extended Data Figure 8

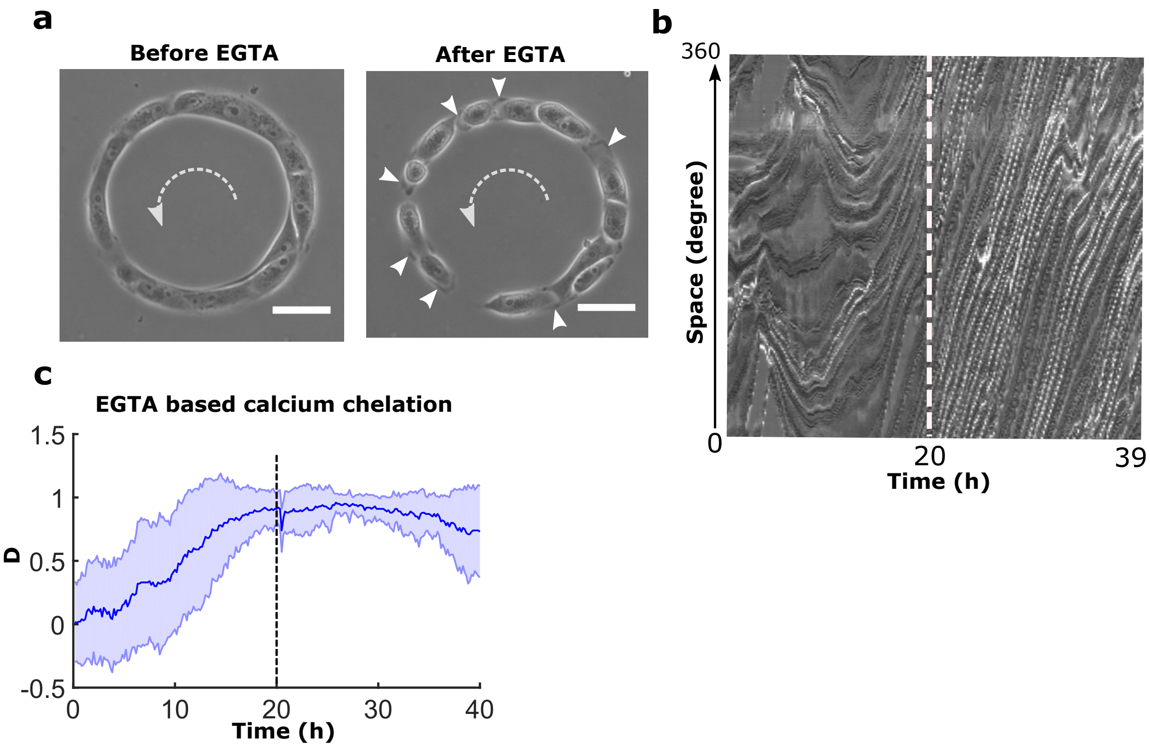

### Extended Data Figure 9

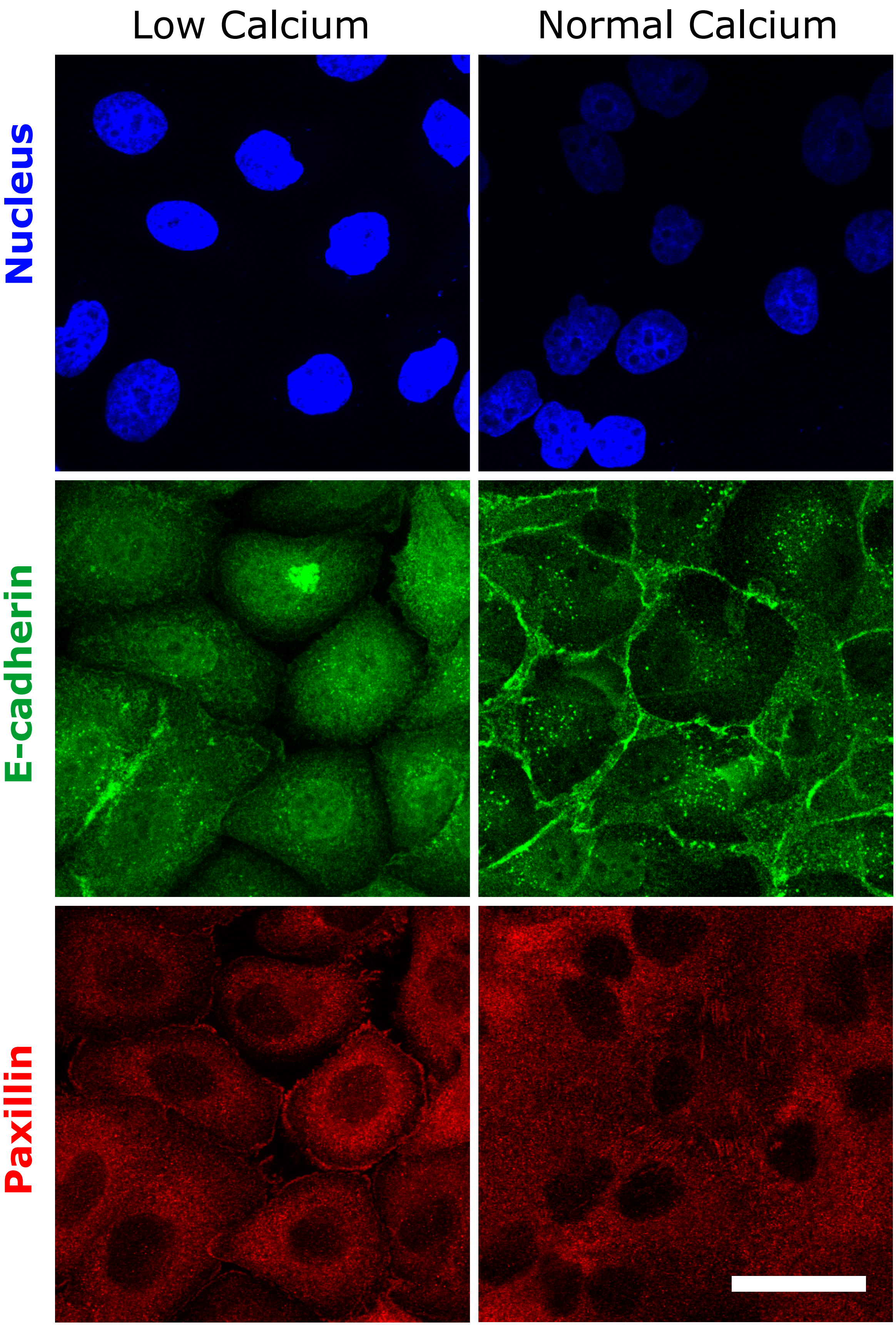

### Extended Data Figure 10

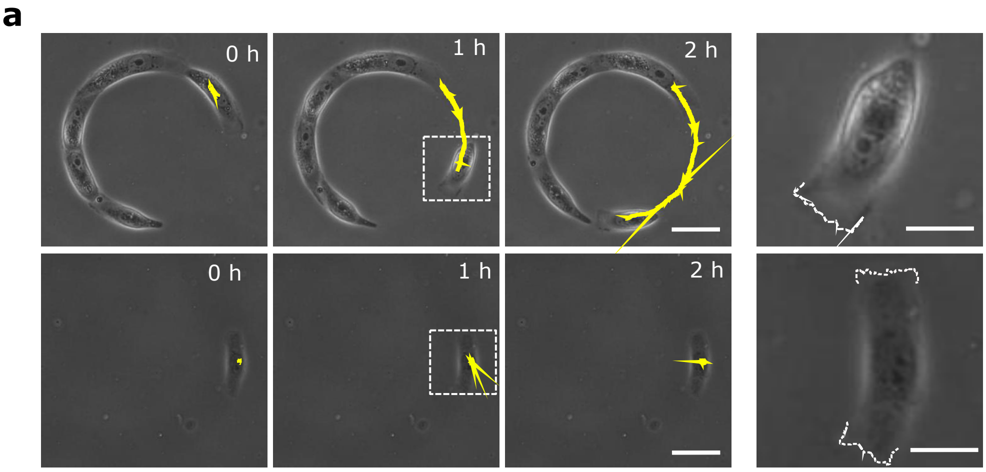


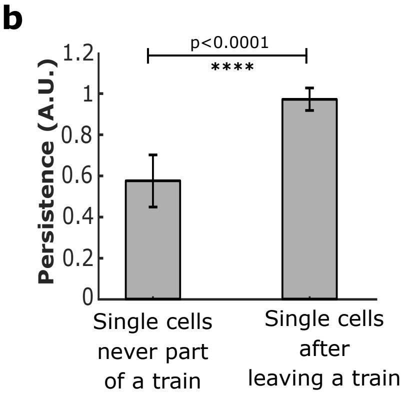

### Extended Data Figure 11

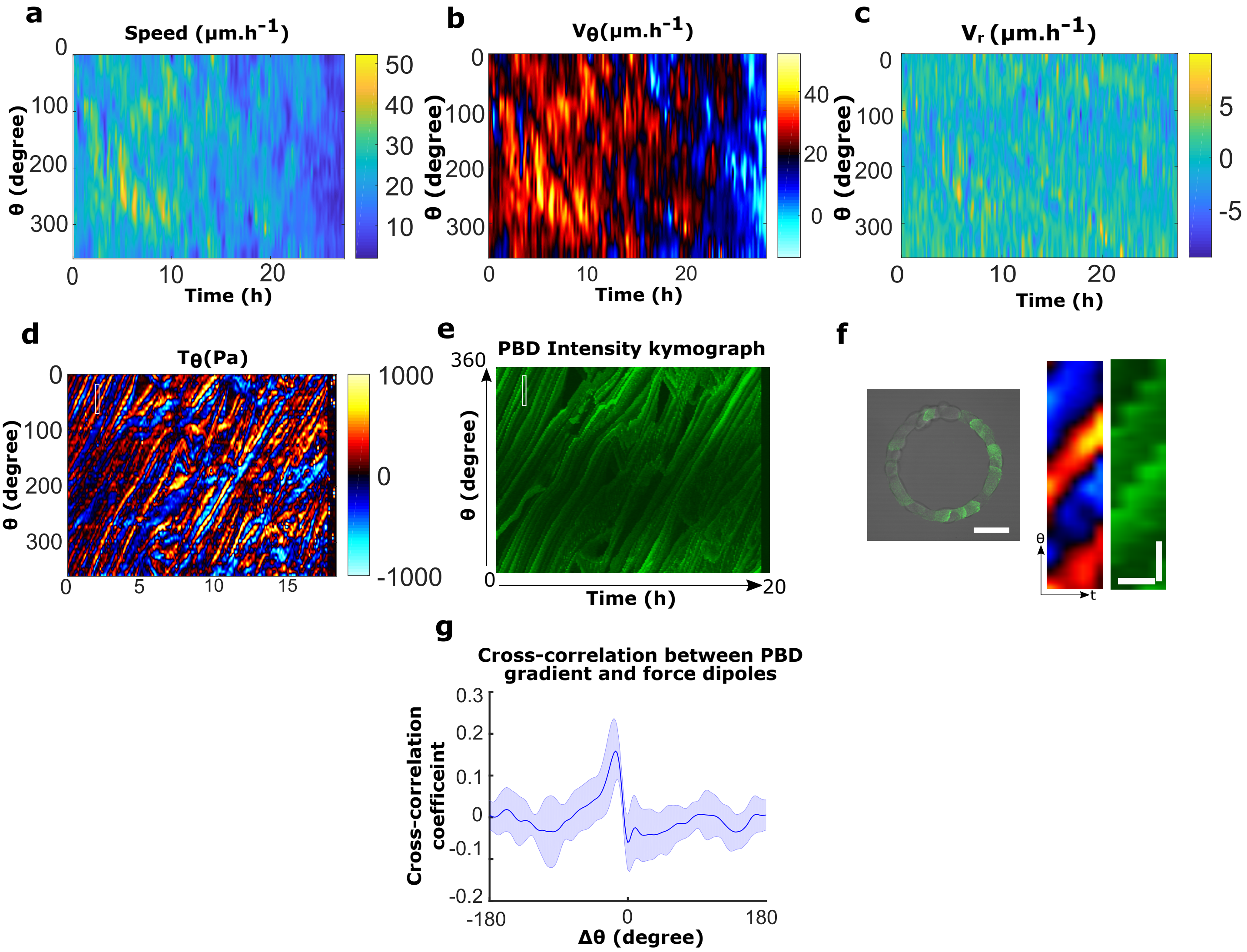

### Extended Data Figure 12

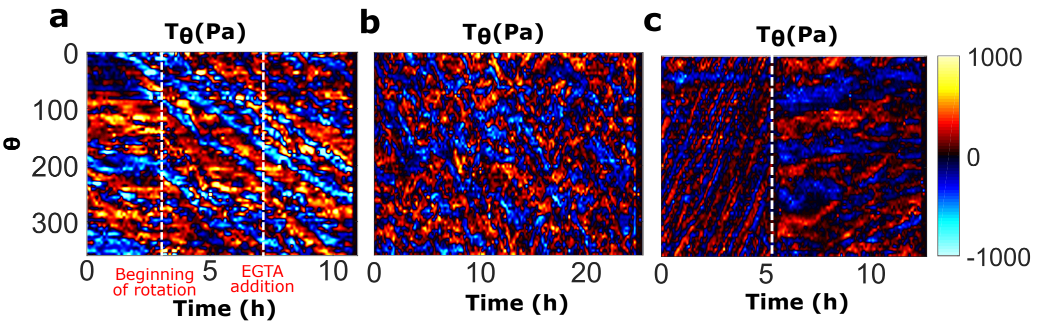

### Extended Data Figure 13

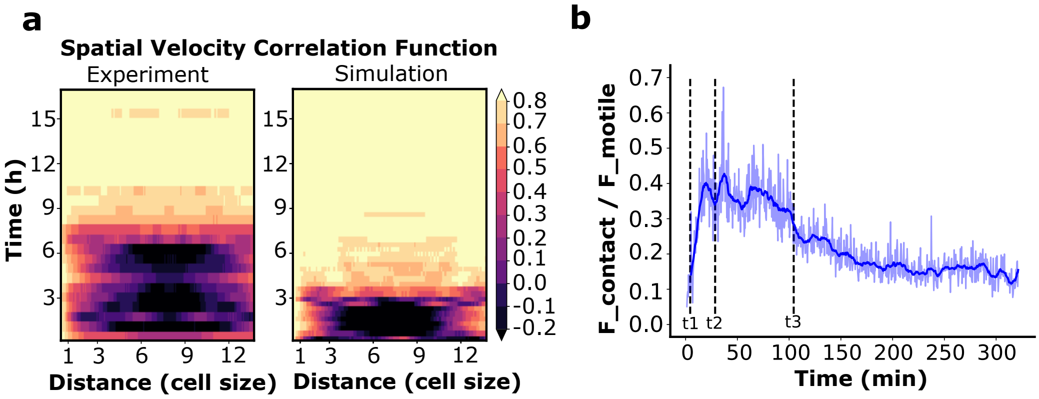

### Extended Data Figure 14

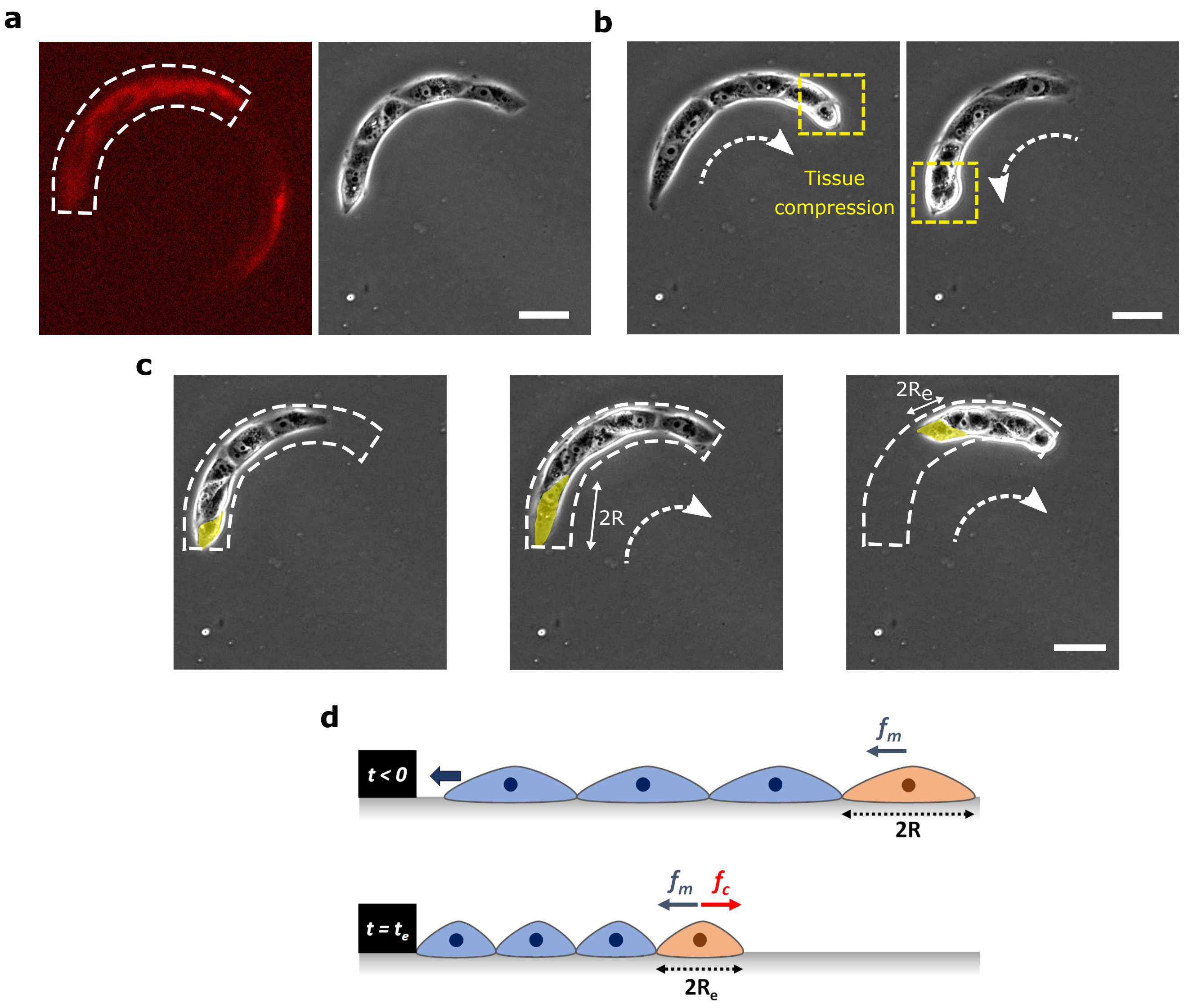

**Supplementary figure legends:**

**Extended Data Figure 1 | Characterization of collective rotation in different conditions and different cell lines**

**a,** Distribution of rotation direction in rings (n=292, m=10). **b, c,** Coordination parameter and rotation speed evolution over time in the rings for cells under normal conditions (no-mitomycin, n=46,m=3) or proliferation blocked (with mitomycin, n=62,m=3) condition. The plateau corresponds to the emergence of a persistent rotation. **d,** Average speed of rotation for mitomycin treated (n=62, m=3) and untreated cells (n=46, m=3). **e,** Coordination parameters for different cell lines, Eph4 (n=10) and Caco2 (n=30). **f,** Coordination parameters *D* for MDCK α-catenin KD cells (n=30) and MDCK Snail expressing cells (n=30).

**Extended Data Figure 2 | Repolarization time scale of single cell and single cell polarities.**

**a-b,** Spatio-temporal montage of MDCK-PBD-YFP cells showing repolarization of a single cell upon collision. White arrows (dashed) show the direction of cell migration. Red arrow heads pointing at time indicate the time of the collision (at 2 min.) and the appearance of Rac1 repolarization signal (at 6 min.), respectively. **c,** Confocal image of basal plane in a 50 µm wide ring shows a PBD gradients in all the cells indicating cell scale polarized Rac1 activities. **d,** Line intensity profile (as visualized by PBD biosensor intensity) shows average Rac1 gradient of single cells (n=42 from 2 rings, m=2) from multiple rings (as represented in a single cell marked by line -ab in panel **c). e,** MDCK cells expressing YFP-PBD and mcherry-AHPH. **f,** Saptio-temporal kymograph shows the single cells (marked with white dashed line) retain the front-rear characteristics during rotations. **g,** Distribution of AHPH (red) and PBD (green) at single cell level from the orthogonal view at one time point. Typical example of a single cell with boundaries marked in dashed white line (Scale bar: 2 µm). **h,** Line graph showing the intensity distribution of PBD and AHPH in single cells (n= 5 cells) of a rotating ring. **i,** Schematics showing the gradient of PBD and AHPH in single cells of a train. All scale bars unless mentioned specifically: 50 µm.

**Extended Data Figure 3 | Immunostainings of cell-substrate adhesions (paxillin) and cell-cell adhesion (E-cadherin).**

**a,** Immunofluorescence for E-cadherin-GFP (Z-projection) in a rotating ring. Anti-clockwise direction is indicated by the white dashed arrows. The diffused E-cadherin distribution indicates the cryptic lamellipodia. **b,** Basal immunofluorescent paxillin staining labelled focal adhesions. **c,** Nuclei labelled in blue. **d,** Merge E-cadherin (green), paxillin (green) and nuclei (blue). **(e-f)** Images showing enlarged views of (e'-f'). **g,** Schematic showing single cells in rotating ring with cryptic lamellipodia revealed by polarized distributions of E-cadherin and paxillin Scale bars: 50 µm **(a-d)**, 20 µm **(e-f)**

**Extended Data Figure 4 | Cell confinement induces large protrusive activity and high speed of cellular protrusions at the cell front.**

**a,** Schematics explaining the experimental set-up to confine front cell extensions. **b,** Representative images of actin-GFP cells on fibronectin coated glass substrates. Before confinement (BC) shows cells with a free leading edge and protruding activity. After confinement

(AC) shows cells whose leading edges are confined underneath the confining PDMS block (scale bar: 10 µm). **c,** Kymograph of the cell front before (top) and after confinement (after) (scale bar: 5 µm). **d,** Cell edge position over time before and after confinement. Negative time correspond to the front edge position before confinement and positive time after confinement (n=10). The slope of curve (dashed magenta line) indicates the velocity of cell protrusions before (0.27 µm/min) and after confinement (1.58 µm/min), respectively.

**Extended Data Figure 5 | Measurement of cell polarity using orientation of nucleus-centrosome (NC) axis**.

**a,** Schematic of nuclear-centrosome axis in a migrating polarized cell. The NC axis orientation is determined in cells double stained for nuclei (green) and centrosomes (red). This state of polarity is referred here as 'CN' when centrosome is placed ahead of nucleus in the direction of migration. When centrosome is placed behind the nucleus, then this state is referred as 'NC'. Cells with multiple centrosomes and with centrosome either pointing radially inward or outward of the rings were categorized under "No Orientation". **b,** NC axis orientation in uncoordinated sub-confluent rings (n=141, 13 rings) compared to confluent coordinated rotating rings (n=131, 11 rings). **c-d,** Representative immunofluorescence images of nucleus-centrosome positions in conditions of low calcium before the establishment of a coordinated rotation (n=216,12 rings), normal calcium conditions (n=191, 15 rings) and low calcium treatment after the initiation of coordinated rotation (n=385, 23 rings).

*****Conditions where rings were not rotating persistently in any direction (sub-confluent and low calcium case before rotation), a reference direction is chosen (i.e. clockwise) in order to calculate the alignment of nucleus-centrosome axis. Scale bar: 50 µm

**Extended Data Figure 6 | Effects of Rac1, Arp2/3 and myosin-II inhibitions on rotating cell trains.**

**a,** PBD signal distribution before (top) and after (bottom) addition of the Rac1 inhibitor Z62954982 (100 µM) on rotating rings. **b,** Intensity profile of PBD signal in entire rings before (top) and after (bottom) drug treatment. Expanded views (marked in red box) showing the disappearance of a single cell PBD signal gradient after drug addition. **c,** Kymograph of the PBD images traced along the circumferential midline of the ring track; the white dashed line indicates drug addition. **d, e,** Coordination parameters and speeds measured before and after Rac1 inhibition. Black dashed lines indicate drug addition (n=5, m=2). **f,** Coordination parameters evolution before and following Arp2/3 inhibition (CK666 ,100 µM, n=37) and myosin-II inhibition (Blebbistatin, 80 µM, n=30). Dashed vertical lines (blue and red color) indicate the point of drug addition respectively. All scale bars: 50 µm.

**Extended Data Figure 7 | Rac1 polarity gradient perturbation using optogenetics.**

**a-b,** Time evolution of PBD signal in cells after homogeneous photoactivation of TIAM. Post activation, TIAM begins to localize at cell membrane. Within 5 minutes of the photoactivation, the PBD gradient flattens in cell indicating the loss of Rac1 polarity in the cell (n=11, from 7 cells). Intensity is measured along the dashed yellow line as indicated in representative panel **a.** Scale bar: 20 µm.

**Extended Data Figure 8 | Compromising cell-cell adhesion strength by EGTA treatment.**
**a,** Phase contrast imaging shows the loosening of cell-cell junctions after EGTA treatment. Exposed cryptic lamellipodia of migrating cells are marked by white arrowheads after cell-cell junction loosening. The dashed white arrows indicate the direction of rotation. **b,** Representative displacement kymograph of phase contrast imaged cells traced along the circumferential midline of the ring track before and after EGTA treatment (2 mM) (white dashed line indicates the time of EGTA addition ~ 20 h). **c,** Coordination parameter of rotating cell rings, before and after EGTA treatment (black dashed line indicates the time of EGTA addition, 20 h) (n=40, m=1). Scale bars: 50 µm.

**Extended Data Figure 9 | Cell-cell junctions and focal adhesions in the presence of low and normal calcium media.**
Nucleus (blue)-E-cadherin (green)-paxillin (red) immunofluorescence of cell monolayers. E-cadherin accumulation at cell-cell contacts detected in normal medium, disappear in low calcium. By contrast, paxillin staining reveals the presence of focal adhesions in both conditions. Scale bar: 20 µm.

**Extended Data Figure 10 | Single cell migration persistency in the absence of cell-cell junctions.**
**a,** Top panel: Phase contrast images of cell trains at low density that show spontaneous detachment of single cells at the free edges. Sequential images showing cell detachment followed by a polarized persistent migration. Expanded view of the dashed white box reveals the active lamellipodial activity on one side of the cell (marked with white dashed line). Bottom panel: Control experiment showing the behavior of a single MDCK cell with no previous contact with other cells. No preferential migration is observed. Both edges of the cell show lamellipodial activity (marked by dashed white line in the expanded view of white dashed box; Scale bars: 20 µm). The yellow lines indicate the distance travelled by the cells at given times. **b,** Persistence of cell movements for single isolated cells (n=12) and for single cells detaching from sub-confluent trains (n= 11). All scale bars unless mentioned specifically: 50 µm.

**Extended Data Figure 11 | Spatio-temporal distributions of velocity profiles and tangential traction forces correlated with PBD profiles during rotation.**
**a,** Spatio-temporal kymograph of cell train speed for the experiment referred in Fig.2c. **b,** Spatio-temporal tangential speed kymograph. **c,** Spatio-temporal radial speed kymograph. **d,** Tangential traction force spatio-temporal profile of MDCK-PBD cells in rotating ring. **e,** Corresponding PBD intensity profile and tangential traction force distribution as shown in panel **d**. **f,** MDCK-PBD cells on ring pattern (Left). Expanded view of corresponding regions of interest for PBD gradients and traction force dipoles (marked by white rectangular box in panel d, e) (right). Scale bar: 10 µm. (ϴ), 30 minutes (t). **g,** Cross correlation of tangential traction force profile and PBD signal show a high spatial correlation (n=10,m=2). All scale bars unless mentioned specifically: 50 µm.

**Extended Data Figure 12 | Traction force distribution in α-catenin KD cells, Arp2/3 inhibition and under EGTA induced low calcium condition**

**a,** Tangential traction forces post EGTA treatment indicate single cell level force dipoles in a rotating ring before and after EGTA treatment (beginning of rotation is marked by the white dashed line ~ at 3h), EGTA addition is marked by the white dashed line at ~7h. **b,** Tangential traction force distribution in α-catenin KD cells. **c,** Tangential traction force pattern before and after Apr2/3 inhibition. CK666 drug addition (100 µM) is marked by the white dashed line.

**Extended Data Figure 13 | Simulation for photo-activation, spatial velocity correlation and force dipole transition**

**a,** Spatial velocity correlation *C(x',t)* as a function of distance x' in experiments and simulations (both with 15 cells) showed similar migration dynamics. $C(x',t) = \frac{<u(x+x',t)\times u(x,t)>_x}{\sqrt{<u(x+x',t)^2>_x \times <u(x,t)^2>_x}}$, where x, x' are curvilinear abscissas, u is the angular velocity (positive in counter-clockwise direction) and t is time. **b,** Ratio of contact and motile forces in 1 simulation (taken from Fig. 4b), recapitulating the force-distribution transition observed in Fig. 2f-h: before polarization starts (t<t1), and once collective rotation is established (t>t3), cells migrate as single dipole and their viscous interaction with the substrate is mainly balanced by their motility. During the coordination process (between t1 and t3), cells form larger dipoles and the contact forces contribute to viscous force balance (t1: cells start polarizing, until they are all polarized at t2, and rotate collectively from t3) (blue thick line: smoothing over time).

**Extended Data Figure 14 | Cell compressibility measurement in experiments of polarized cell trains migrating between broken fibronectin micropatterns.**

**a,** Broken ring micropatterns and cell train. **b,** Train migrating towards the end of fibronectin confinement: train at the point of maximal compression. **c,** Simulation parameter measurement of the resting radius *R* and equilibrium radius $R_{e.}$ of the last cell before and after compression as represented by a yellow shaded region. **d,** Schematic: force balance for the last cell of the train. Scale bars: 50 µm.

**Movie Captions:**

**Movie 1:** MDCK cells break symmetry and rotate collectively on 1-D ring confinement. Ring outer diameter 200 µm, track width of 20 µm.

**Movie 2:** MDCK cell rotations under geometrical confinements with different shape (g1-triangle, g2-ellipse, g3-U shape), length(diameter d1:outer-400 µm, middle-200 µm, inner-100 µm, d2-1mm ) and width (w1-50 µm, w2-100 µm, w3-200 µm).

**Movie 3:** Other cell types display collective behavior: **a,** Eph4 and **b,** Caco2

**Movie 4:** Time scale of Rac1 based repolarization in single cells upon collision with the larger train

**Movie 5:** Rac1 based front-rear polarity is established in each cell of the rotating ring after symmetry breaking.

**Movie 6:** Optogenetics: photoactivation of TIAM at 120 minutes in a quarter region of rotating ring (marked by blue dashed line) is indicated by its junctional localization, which follows a gradual arrest of coordinated rotations in ring.

**Movie 7:** MDCK cells fail to initiate rotation and break symmetry **a,** MDCK α-Catenin stable knocked cells and, **b,** MDCK-Snail expressing cells

**Movie 8:** MDCK cells fail to initiate rotation in presence of low calcium condition but breaks symmetry and begins to rotate upon addition of normal calcium. However, once symmetry is broken, MDCK cells continue to rotate even when low calcium condition is reintroduced.

**Movie 9:** Epithelia continuity is not required for the maintenance of rotations: Laser ablation of cells in the rotating ring maintains the directed migration of cells.

**Movie 10: a,** Numerical simulations reproduce the symmetry breaking process and polarity establishment in cell rings. Geometrical shapes represent cell centers. Circles: non-polarized cells. Triangles: polarized cells, pointing in their polarity direction. Cell boundaries are not represented, though blue lines indicate the intensity of contact forces on a cell. **b,** Cell swapping upon migrating train collisions in the case of low cell-cell junction strength-based interactions (top) when compared to cell repolarization during collision in normal cell-cell junction strength (bottom).
